# Role of PARP1 in oligodendrocyte differentiation during developmental myelination and remyelination after myelin damage

**DOI:** 10.1101/2021.04.23.441060

**Authors:** Yan Wang, Sheng Zhang, Bokyung Kim, Vanessa L. Hull, Jie Xu, Preeti Prabhu, Maria Gregory, Veronica Martinez-Cerdeno, Xinhua Zhan, Wenbin Deng, Fuzheng Guo

## Abstract

The function of poly(ADP-ribosyl) polymerase 1 (PARP1) in myelination and remyelination of the central nervous system (CNS) remain enigmatic. Here we report that PARP1 is an intrinsic driver for oligodendroglial development and myelination. Genetic PARP1 depletion impairs the differentiation of oligodendrocyte progenitor cells (OPCs) into oligodendrocytes and impedes CNS myelination. Mechanistically, PARP1-mediated PARylation activity is not only necessary but also sufficient for OPC differentiation. At the molecular level, we identify the RNA-binding protein Myef2 as a novel PARylated target which we show controls OPC differentiation through PARylation-modulated de-repression of myelin protein expression. Furthermore, PARP1’s enzymatic activity is necessary for oligodendrocyte and myelin regeneration after demyelination. Together, our findings suggest that PARP1-mediated PARylation activity may be a potential therapeutic target for promoting OPC differentiation and remyelination in neurological disorders characterized by arrested OPC differentiation and remyelination failure such as multiple sclerosis.

## Introduction

Oligodendrocytes (OLs), the myelin-forming cells in the central nervous system (CNS), are differentiated from oligodendrocyte progenitor cells (OPCs). The differentiation of OPCs into OLs and subsequent myelination by OLs are essential for maintaining normal CNS functions. Defects of these two events result in nerve conduction disruption, axonal dysfunction, and neurological disabilities such as in periventricular leukomalacia, multiple sclerosis, and leukodystrophy (Bercury and Macklin, 2015). Hence, understanding the mechanisms underlying OL development and myelination is crucial for devising therapeutic strategies to promote OPC differentiation and myelinogenesis in CNS dysmyelinating and demyelinating diseases.

The nuclear protein poly (ADP-ribose) polymerase 1 (PARP1) is the founding member of the PARP family consisting of at least 17 members which transfer single ADP-ribose unit [mono (ADP-ribose)] or poly (ADP-ribose) units [poly(ADP-ribose), PAR] to target proteins (Gupte et al., 2017). PARP1 is one of the four members (PARP1, PARP2, PARP5a, and PARP5b) that catalyze the covalent attachment of PAR onto target proteins, a process called PARylation (Ke et al., 2019). PAPP1 has been reported responsible for >90% of cellular protein PARylation in non-neural tissues (Kamaletdinova et al., 2019), which can be reversed by the activity of the degrading enzyme PAR glycohydrolase (PARG) (Davidovic et al., 2001). Initially discovered as an essential player for DNA repair in cancer cells, PARP1 has been increasingly recognized as a key regulator for diverse cellular processes such as chromatin remodeling, gene expression regulation, and cell death/survival (Gupte et al., 2017). Despite the growing knowledge of its basic molecular functions, the physiological role and significance of PARP1 in oligodendroglial development and myelinogenesis are poorly defined.

In the developing murine CNS, *Parp1* transcripts are expressed in all neural cells with the highest abundance in oligodendroglial lineage cells (www.brainrnaseq.org) (Zhang et al., 2014). Under demyelinating conditions, PARP1 and/or its PARylation activity has been reported to be upregulated in various neural and immune cells in multiple sclerosis (Farez et al., 2009; Meira et al., 2019; Veto et al., 2010). In the past three decades, the role of PARP1 in regulating neuroinflammation has been extensively studied albeit with descrepant or even conflicting results (Cavone and Chiarugi, 2012). However, little is known about if and how PARP1 regulates oligodendroglial lineage development and CNS myelinogenesis. This is reflected by an extremely limited amount of data in the field of PARP1 and oligodendroglial lineage cells including those from our own group (Baldassarro et al., 2017; Plane et al., 2012).

In the current study, we showed that, in contrast to its role in cell death/survival under genotoxic stress conditions, PARP1 is an intrinsic positive regulator for oligodendrocyte development. By generating and characterizing *Parp1* knockout (KO) and lineage-specific *Parp1* conditional KO (cKO) mice, we unequivically demonstrated that PARP1 is necessary for OPC differentiation and developmental myelination. We found that PARP1’s PARylation activity plays a key role in PARP1-regulated OPC differentiation and myelin gene expression both *in vivo* and *in vitro* in a cell-autonomous manner. At the molecular level, we unveiled that PARP1 PARylates the novel target protein myelin expression factor 2 (Myef2) and disociates it from binding with myelin gene mRNAs, relieving its suppressive effects on myelin protein expression. Thus, our findings reveal a yet unappreciated role of PARP1 in oligodendroglial development and CNS myelination and suggest that augmenting PARP1 and/or its activity-mediated PARylation might be a viable option to enhance OPC differentiation and remyelination in demyelinating disorders.

## Results

### PARP1 is dynamically expressed in OL lineage during developmental myelination

We previously reported that the stem cell factor Sox2 promotes oligodendrogenesis during CNS developmental myelination (Zhang et al., 2018a; Zhang et al., 2018b). We therefore sought to screen Sox2 interacting partners at the proteomic level using co-immunoprecipitation (Co-IP) followed by liquid chromatography and tandem mass spectrometry identification (LC-MS/MS) (fig 1A). This effort identified PARP1 as one of the top candidates of Sox2’s interactome (fig. S1B-C) in primary OLs. Co-IP/Western blot (fig. S1D) further confirmed the interaction between PARP1 and Sox2 in OLs. Triple immunohistochemistry (IHC) showed that PARP1 and Sox2 were co-expressed in Sox10^+^ oligodendroglial lineage cells *in vivo* (fig. S1E).

The expression dynamics of PARP1 in the oligodendroglial lineage cells (OLCs) has yet to be defined. Western blotting assay demonstrated that PARP1 was highly expressed during developmental myelination and progressively downregulated after myelination completion in the murine CNS (Fig. 1A). Within the OLCs, triple IHC showed that PARP1 was expressed at a relative low level in PDGFRα^+^ OPCs (Fig. 1B, arrowheads), upregulated in CC1^+^ premyelinating OLs (Fig. 1B, arrows) which were identified by a premyelinating OL marker TCF7l2 (Fig. 1C, arrowheads) (Hammond et al., 2015; Marques et al., 2016), and downregulated in fully mature myelinating OLs in the adult spinal cord where PARP1 was maintained at low level in PDGFRα^+^ OPCs (Fig. 1D). Using primary OPC cultures purified from neonatal mouse brain (Lang et al., 2013; Zhang et al., 2018b), we found that PARP1 mRNA (Fig. 1E) and protein (Fig. 1F) level was low in OPCs, transiently upregulated in newly differentiated OLs, and downregulated in mature OLs. This dynamic expression among OL lineage progression and maturation (Fig. 1G) suggests that PARP1 may regulate oligodendroglial lineage progression during postnatal development.

**Fig. 1.**
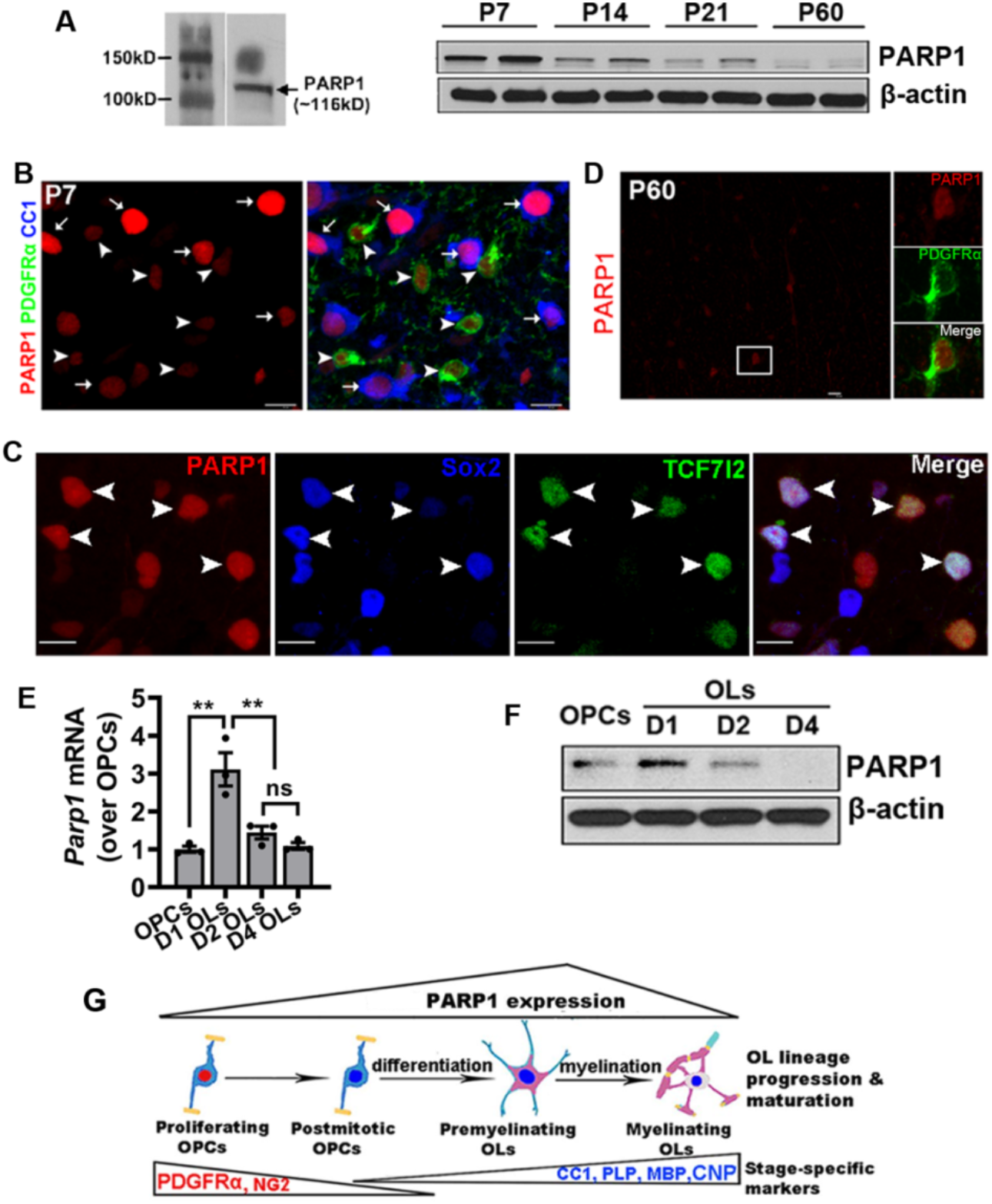
PARP1 expression along oligodendrocyte lineage progression and maturation. **A,** Western blotting assay for PARP1 (∼116kD) in the spinal cord at different time-points (P, postnatal day). *β*-actin was used as an internal loading control. **B,** triple IHC of PARP1, OPC marker PDGFR*α*, and OL marker CC1 in P7 spinal cord. Arrowheads, PARP1^+^/PDGFRα^+^ OPCs, arrows, PARP1^+^/CC1^+^ OLs. **C,** representative confocal images of PARP1, Sox2, and TCF7l2 in P7 spinal cord. Arrowheads, PARP1^+^/SOX2^+^ cells were positive for TCF7l2^+^ (arrowheads), a marker of newly differentiated OLs. **D,** IHC of PARP1 in adult spinal cord. Boxed area shown at the right. **E-F,** PARP1 mRNA (**E**) and protein (**F**) in brain-derived primary OPCs and differentiated OLs at day 1, 2, and 4. D1 and D2 OLs, newly differentiated OLs; D4 OLs, fully matured OLs. One-way ANOVA followed by Tukey’s multiple comparison test, *F*_(3,8)_=16.63, *P* = 0.0008; ** *P* < 0.01. N = 3 replicates each group. **G** Schematic drawing depicting the temporal dynamics of PARP1 during OL development: low expression in OPCs, upregulation in newly differentiated, premyelinating OLs, and downregulation in mature, myelinating OLs. Scale bars: B, C, D, 10μm.

### PARP1 deficiency results in impaired OL maturation and developmental myelination

PARP1 is well-known as a key mediator for DNA damage response and repair (Pascal, 2018). Little is known about its role in CNS myelination. To elucidate the physiological importance of PARP1 in developmental myelination, we first characterized and compared oligodendrogenesis and myelin formation in the CNS of *Parp1* KO mice (Plane et al., 2012; Wang et al., 1995) with their wild type littermates (*Parp1* WT). Western blotting and qRT-PCR confirmed the absence of PARP1 protein (Fig. 2A) and mRNA (Fig. 2B) in *Parp1* KO mice. We observed prominent hypomyelination in the brain of P10 *Parp1* KO mice compared with WT littermates (Fig. 2C-D). The density of Sox10^+^CC1^+^ differentiated OLs but not Sox10^+^PDGFRα^+^ OPCs was significantly diminished in the corpus callosum of P10 *Parp1* KO compared with WT littermates (Fig. 2E-F), suggesting that PARP1 regulates oligodendroglial maturation. To corroborate this conclusion, we found a marked decrease in the brain mRNA levels of major myelin proteins and comparable mRNA level of PDGFRα between *Parp1* KO and WT littermates (Fig. 2G). Previous studies show that the nuclear protein TCF7l2 is transiently expressed in newly formed premyelinating OLs (Hammond et al., 2015; Marques et al., 2016; Zhao et al., 2016). Therefore, the number of TCF7l2^+^Sox10^+^ cells at any given time provides a quantitative readout for the rate of oligodendrogenesis. We observed a significant diminution of TCF7l2^+^Sox10^+^ cell density in the corpus callosum of P10 *Parp1* KO mice (Fig. 2H-I). Consistently, the expression of other premyelinating OL-enriched marker genes *Bmp4, Ugt8a, Bcas1, and Gpr17*, was all significantly reduced in P10 *Parp1* KO brain (Fig. 2J). These data suggest that PARP1 deficiency inhibits oligodendrocyte maturation and developmental myelination.

**Fig. 2.**
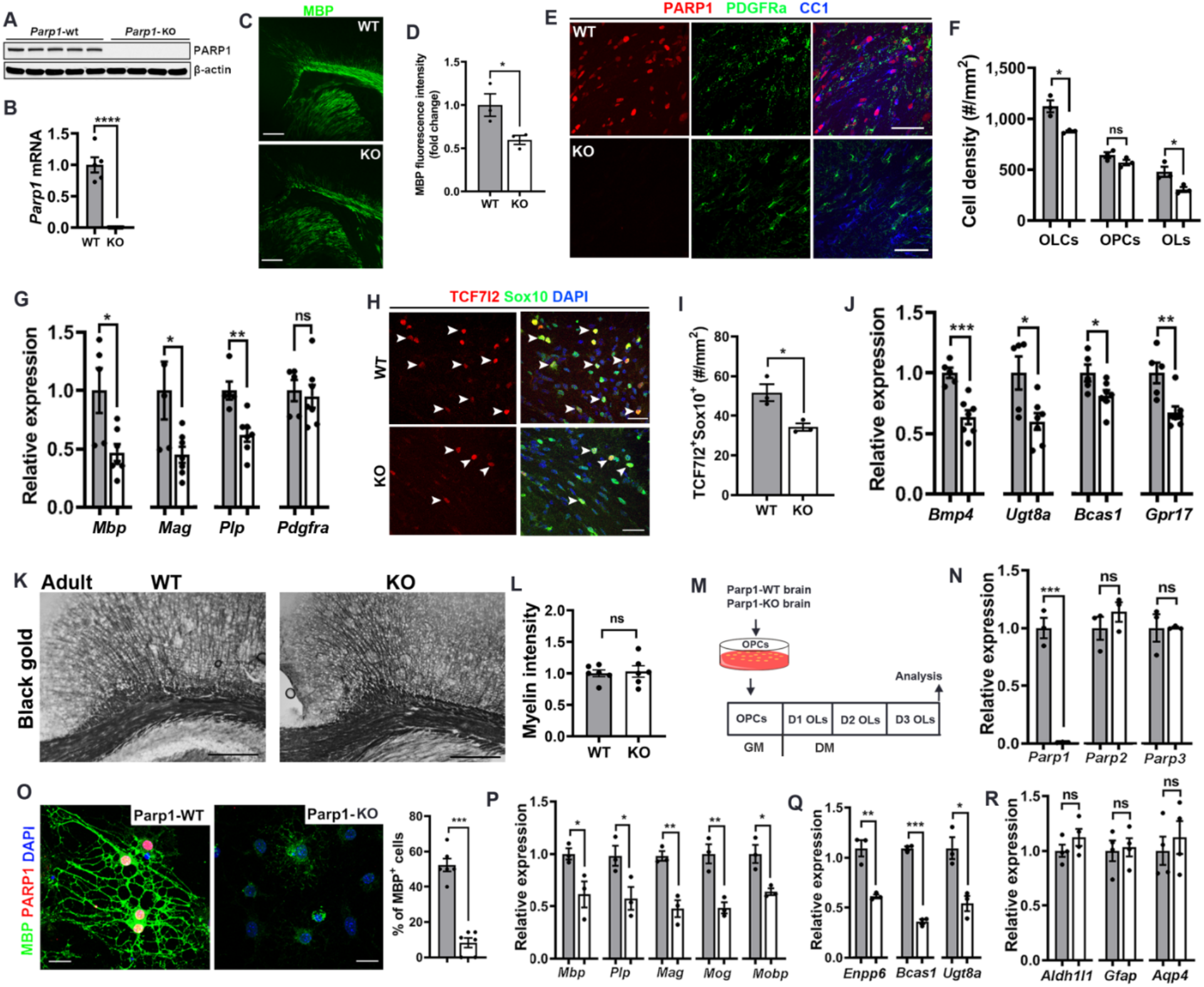
*Parp1* deficiency inhibits OL maturation and results in hypomyelination during postnatal development. **A,** Western blotting and quantification of P7 spinal cord protein from WT and *Parp1* KO mice. N = 5 WT, 4 *Parp1* KO. **B,** qRT-PCR assay (using primers amplifying *Parp1* exon 2 mRNA) for *Parp1* transcripts in P7 spinal cord. N = 5 WT, 7 *Parp1* KO, *t*_(10)_ = 9.926 *P* < 0.0001. **C-D,** representative confocal images for myelin stained by myelin basic protein MBP (**C**) and relative intensity (**D**) in P10 forebrain. N = 3 each group, *t*_(4)_ = 2.895 *P* = 0.0443. **E**, representative confocal images of PARP1, PDGFRα, and CC1 in P10 corpus callosum. **F**, densities of Sox10^+^ OLCs, Sox10^+^PDGFRα^+^ OPCs, and Sox10^+^CC1^+^ OLs in the corpus callosum of P10 mice. N = 3 WT, 3 *Parp1* KO. OLCs, *t*_(4)_ = 4.182 *P* = 0.0139; OPCs, *t*_(4)_ = 1.829 *P* = 0.1414; OLs, *t*_(4)_ = 3.117 *P* = 0.0356. **G,** qRT-PCR assay for myelin-specific genes and *Pdgfra* in the brain. N = 5 WT, 7 *Parp1* KO. *Mbp*, *t*_(10)_ = 2.932 *P* = 0.0150; *Mag*, *t*_(10)_ = 2.483 *P* = 0.0324; *Plp*, *t*_(10)_ = 3.923 *P* = 0.0029; *Pdgfra*, *t*_(10)_ = 0.3517 *P* = 0.7323. **H-I,** representative confocal images showing TCF7l2^+^Sox10^+^ newly formed premyelinating OLs (**H**, arrowheads) and quantification (**I**) in the corpus callosum of P10 mice. N = 3 WT, 3 *Parp1* KO, *t*_(4)_ = 3.733 *P* = 0.0202. **J**, qRT-PCR assay for genes encoding pre-myelination OL markers in the brain. N = 5 WT, 7 Parp1 KO. *Bmp4*, *t*_(10)_ = 4.775 *P* = 0.0008; *Ugt8a*, *t*_(10)_ = 2.855 *P* = 0.0171; *Bcas1*, *t*_(10)_ = 2.364 *P* = 0.0397; *Gpr17*, *t*_(10)_ = 3.487 *P* = 0.0059. **K-L**, Black Gold II myelin staining (**K**) and quantification (**L**) showing comparable myelination in the corpus callosum and the overlaying cingulate cortex between *Parp1* KO (n = 6) and WT mice (n = 6) at P70. *t*_(10)_ = 9.705 *P* < 0.0001. **M**, experimental design for Panel **N-R**. GM, growth medium, DM, differentiation medium. **N**, qRT-PCR assay confirming depletion of *Parp1* and non-perturbation of *Parp2* and *Parp3*. N = 3 WT, 3 *Parp1* KO. *Parp1*, *t*_(4)_ = 11.22 *P* = 0.0004; *Parp2*, *t*_(4)_ = 1.079 *P* = 0.3415; *Parp3*, Welch-corrected *t*_(2.019)_ = 0.0260 *P* = 0.9816. **O**, representative confocal images of MBP and PARP1 and quantification of MBP^+^ ramified cells among DAPI^+^ cells. N = 6 WT, 6 *Parp1* KO, *t*_(10)_ = 9.705 *P* < 0.0001. **P-Q**, relative expression of myelin-specific genes (**P**) and pre-myelinating OL marker genes (**Q**) quantified by qRT-PCR (n = 3 WT, 3 *Parp1* KO). *Mbp*, *t*_(4)_ = 2.867 *P* = 0.0456; *Plp*, *t*_(4)_ = 2.794 *P* = 0.0491; *Mag*, *t*_(4)_ = 5.412 *P* = 0.0056; *Mog*, *t*_(4)_ = 4.860 *P* = 0.0083; *Mobp*, *t*_(4)_ = 3.909 *P* = 0.0174; *Enpp6*, *t*_(4)_ = 5.738 *P* = 0.0046; *Bcas1*, *t*_(4)_ = 24.70 *P* = <0.0001; *Ugt8a*, *t*_(4)_ = 4.149 *P* = 0.0143. **R**, qRT-PCR assay for astrocyte-specific markers (n = 4 WT, 4 *Parp1* KO). *Aldh1l1*, *t*_(6)_ = 1.268 *P* = 0.2517; *Gfap*, *t*_(6)_ = 10.2500 *P* = 0.8109; *Aqp4*, *t*_(6)_ = 0.6142 *P* = 0.5617. Unpaired, two-tailed Student’s *t* test, \**p* < 0.05; \*\**p* < 0.01; \*\*\**p* < 0.001, ns, not significant. Scale bars: C, 200μm; E, H, 50μm; K, 100μm; O, 10μm. Gray bars: WT, white bars: *Parp1* KO.

We next analyzed the adult brain of *Parp1* KO mice to interrogate if PARP1 is required for myelin maintenance. Brain myelination, visualized by Black Gold II myelin staining (Fig. 2K), was comparable between *Parp1* KO and WT mice (Fig. 2L) at P70. Consistently, no significant differences were detected in the densities of Sox10^+^CC1^+^ mature OLs, Sox10^+^PDGFRα^+^ OPCs, and Sox10^+^ OLCs (fig. S2A) in the adult corpus callosum (fig. S2B) and the cortex (fig. S2C), nor in the brain mRNA levels of major myelin proteins and premyelinating OL marker TCF7l2 (fig. S2D). Intriguingly, Rotarod test (Hull et al., 2020; Zhang et al., 2018b) demonstrated that the motor function of adult *Parp1* KO mice was markedly impaired, as evidenced by significantly decreased time latency to fall from rod (fig. S2E-F).

### PARP1 is a cell intrinsic regulator for oligodendrocyte maturation

To determine if PARP1 regulates oligodendrocyte maturation in a cell autonomous manner, we purified primary OPCs from neonatal *Parp1* KO and WT brain and induced OPC differentiation and maturation by chemical-defined differentiation medium (DM) in the dish (Fig. 2M). PARP1 depletion did not affect the expression of the family members PAPR2 and PARP3 (Fig. 2N). Strikingly, when cultured in the DM for 3 days, PARP1-depleted OPCs exhibited severe inhibition of differentiation and maturation, as shown by reduced density of MBP^+^ OLs with arborized morphology (Fig. 2O), diminished expression of major myelin proteins (Fig. 2P), and decreased level of pre-myelinating OL markers (Fig. 2Q) compared with those in WT cultures. We observed no difference in the expression of astrocyte-specific genes *Aldh1l1, Gfap*, and *Aqp4* (Fig. 2R), suggesting PARP1-depleted OPCs do not switch their fate to astroglial lineage. Collectively, our data indicate that PARP1 regulates oligodendroglial differentiation and maturation in a cell autonomous manner.

### PARP1 cKO in Olig2-expressing cells inhibits oligodendrogenesis and developmental myelination

In addition to oligodendroglial lineage, PARP1 is also expressed in other neural cells of the CNS (http://www.brainrnaseq.org/) (Zhang et al., 2014) and in non-neural tissues. Therefore, we employed *Cre-loxP*-based conditional knockout (cKO) strategy to elucidate PARP1’s role in oligodendroglial development. To this end, we crossed *Olig2-Cre* (Schuller et al., 2008) with *Parp1-*floxed (Luo et al., 2017) mice to deplete PARP1 selectively in Olig2-expressing primitive OPCs starting from the mid-embryonic stages (Rowitch, 2004). Our fate-mapping study (fig. S3A) demonstrated that *Olig2-Cre*-mediated recombination was observed in up to ∼80% of Sox10^+^ oligodendroglial lineage cells in the corpus callosum of P17 *Olig2-Cre:Rosa26-EYFP* mice (fig. S3B-C), which is in line with previous data (Chavali et al., 2020; Fancy et al., 2014). *Parp1* cKO (fig. S3D) resulted in hypomyelination (fig. S3E-F) and defective oligodendroglial maturation (fig. S3G-H) in the brain of P14 *Olig2-Cre:Parp1* cKO mice compared with Ctrl littermates. However, brain myelination (fig. S3I-J) and mature OL number (fig. S3K) were comparable between *Olig2-Cre:Parp1* cKO and control mice in the adult ages, similar observations to those in adult *Parp1* KO mice. Despite the comparable oligodendroglial phenotypes, adult *Olig2-Cre:Parp1* cKO mice displayed motor function impairment, as evaluated by accelerating Rotarod test (fig. S3L-M). Collectively, our data suggest that PARP1 expression in Olig2-expressing cells is required for developmental myelination in early postnatal mice and motor function as well in adult mice.

### PARP1 regulates OPC differentiation into OLs in the postnatal CNS

Oligodendrocytes in the postnatal corpus callosum and cortex are differentiated from OPCs that are generated from cortical progenitors after birth in the murine brain (Kessaris et al., 2006). To determine the role of PARP1 in postnatal OPC differentiation, we generated *Pdgfra-CreER^T2^*:*Parp1*^fl/fl^ (*Pdgfra:Parp1* cKO) mutants, thus enabling time-conditional depletion of PARP1 in OPCs by tamoxifen treatment (Fig. 3A-B). Tamoxifen administration at P1, P2, and P3 resulted in PARP1 depletion in ∼ 90% PDGFRα^+^ OPCs in the corpus callosum (Fig. 3C), which is congruent with our previous studies using the *Pdgfra-CreER^T2^* line (Zhang et al., 2020a; Zhang et al., 2018b). Unbiased bioinformatic GO analysis of bulk brain RNA-seq showed that OPC differentiation and myelination were significantly enriched among the downregulated genes in *Pdgfra:Parp1* cKO brain (Fig. 3D), indicating a defective oligodendroglial development. qRT-PCR demonstrated that the expression of major myelin-specific genes (Fig. 3E) and premyelinating OL-enriched genes (Fig. 3F) was severely affected in *Pdgfra:Parp1* cKO brain. Histological quantification showed a significant reduction in the number of Sox10^+^CC1^+^ differentiated OLs (Fig. 3G) and TCF7l2^+^ premyelinating OLs (Fig. 3H) in *Pdgfra:Parp1* cKO brain compared with littermate controls. Consistently, MBP expression was observed at significantly lower levels in the subcortical white matter of *Pdgfra:Parp1* cKO brain than that of *Parp1* controls whereas SMI312^+^ axonal bundles were comparable between the two groups (Fig. 3I). We observed that OPC-specific PARP1 depletion did not affect oligodendroglial proliferation (fig. S4A-B), cell death (fig. S4C-D), and microglial (fig. S4E-G) and astroglial (fig. S4G-J) activation. These data suggest that PARP1 regulates OPC differentiation in the postnatal murine brain.

**Fig. 3.**
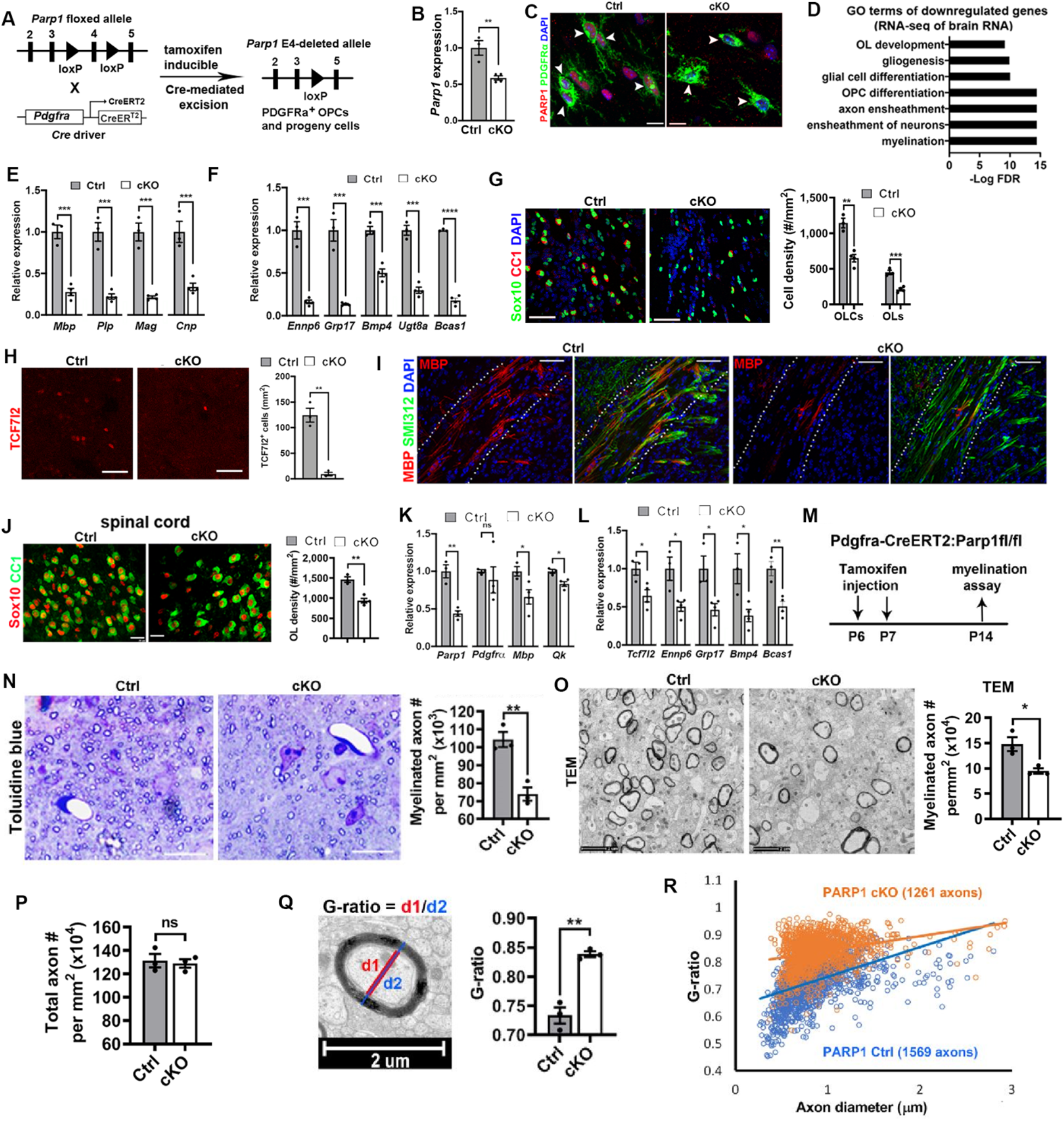
OPC-specific PARP1 cKO inhibits OPC differentiation and myelination. **A,** *Pdgfrα-CreER*^T2^:*Parp1*^fl/fl^ (*Pdgfrα:Parp1* cKO) mice and *Parp1* Ctrl littermate carrying *Parp1*^fl/fl^ or *Pdgfrα-CreER*^T2^ (pooled) were treated with tamoxifen subcutaneously at P1, P2, and P3 (once a day) and analyzed at P9 according to our previous protocols (Zhang et al., 2020a; Zhang et al., 2018b). **B,** qRT-PCR quantification of exon 4-containing *Parp1* mRNA in the brain (n = 3 Ctrl, 4 cKO) *t*_(5)_ = 4.551 *P* = 0.0061. **C**, confocal images showing PARP1 depletion in PDGFRα^+^ OPCs (arrowheads) in the corpus callosum. **D**, gene ontology (GO) analysis showing top enriched biological processes among significantly downregulated genes (cKO vs Ctrl) in the brain (n = 3 Ctrl, 3 cKO). **E-F**, qRT-PCR assays for genes coding myelin-specific proteins (**E**) and pre-myelinating OL markers (**F**) in the brain. N = 3 WT, 4 cKO. *Mbp*, *t*_(5)_ = 8542 *P* = 0.0004; *Plp*, *t*_(5)_ = 7.760 *P* = 0.0006; *Mag*, *t*_(5)_ = 8.591 *P* = 0.0004; *Cnp*, *t*_(5)_ = 5.561 *P* = 0.0026; *Enpp6*, *t*_(5)_ = 9.518 *P* = 0.0002; *Gpr17, t*_(5)_ = 8.181 *P* = 0.0004; *Bmp4*, *t*_(5)_ = 7.603 *P* = 0.0006; *Ugt8a*, *t*_(5)_ = 10.87 *P* = 0.0001; *Bcas1*, *t*_(5)_ = 24.89 *P* < 0.0001. **G**, representative confocal images and densities of Sox10^+^ OLCs and Sox10^+^CC1^+^ OLs in the corpus callosum. N = 3 Ctrl, 4 cKO. Sox10^+^, *t*_(5)_ = 5.965 *P* = 0.0019; Sox10^+^CC1^+^ OLs, *t*_(5)_ = 7.121 *P* = 0.0008. **H**, representative and quantification of TCF7l2-expressing premyelinating OLs in the corpus callosum. N = 3 Ctrl, 3 cKO. TCF7l2^+^, *t*_(4)_ = 8.254 *P* = 0.0012. **I**, IHC of MBP and pan-axonal marker SMI312 in the external capsule (dotted areas). **J**, representative confocal images of Sox10 and CC1 in the ventral white matter of the spinal cord and densities of Sox10^+^CC1^+^ OLs. N = 3 Ctrl, 4 cKO. OLs, *t*_(5)_ = 5.389 *P* = 0.0030. **K-L**, qRT-PCR assays for mRNA expression of *Parp1*, *Mbp*, OPC marker *Pdgfra* and OL marker *Qk* (**K**) and premyelinating OL markers (**L**) in the spinal cord. N = 3 Ctrl, 4 cKO. *Parp1*, *t*_(5)_ = 6.673 *P* = 0.0011; *Pdgfra*, Welch-corrected *t*_(3.096)_ = 0.6521 *P* = 0.5595; *Mbp*, *t*_(5)_ = 2.726 *P* = 0.0415; *Qk*, *t*_(5)_ = 3.950 *P* = 0.0109; premyelinating OL markers, *Tcf7l2*, *t*_(5)_ = 3.002 *P* = 0.0300; *Enpp6*, *t*_(5)_ = 3.394 *P* = 0.0194; *Gpr17*, *t*_(5)_ = 3.203 *P* = 0.0239; *Bmp4*, *t*_(5)_ = 3.253 *P* = 0.0226; *Bcas1*, *t*_(5)_ = 4.482 *P* = 0.0065. **M**, experimental design for **N-R**. The corticospinal tract (CST), a representative white matter tract, was used for analysis. **N**, representative bright field images of toluidine blue myelin staining performed on semithin (500nm) resin sections and quantification of CST axons surrounded by toluidine blue positive myelin. N = 3 Ctrl, 3 cKO. *t*_(4)_ = 5.414 *P* = 0.0056. Scale bar = 10µm. **O**, representative transmission electron microscopic (TEM) images and density of myelinated axons with diameter ≥ 0.3µm. N = 3 Ctrl, 3 cKO. *t*_(4)_ = 3.580 *P* = 0.0232. **P**, density of myelinated and unmyelinated axons (diameter ≥ 0.3μm) in the CST. N = 3 Ctrl, 3 cKO. *t*_(4)_ = 0.3473, *P* = 0.7458. **Q**, diagram and quantification of average G-ratio of myelinated axons in the CST. N = 3 Ctrl, 3 cKO. *t*_(4)_ = 7.170 *P* = 0.0020. **R**, plot of individual G-ratio vs the diameter of each axon. Unpaired, two-tailed Student’s t test, \**P* < 0.05; \*\**P* < 0.01; \*\*\**P* < 0.001, ns, not significant. Scale bars: O, 2μm; C, J, N, 10μm; G, H, I, 50μm.

To determine whether PARP1-regulated OPC differentiation also occurs in the spinal cord, we quantified the number of Sox10^+^CC1^+^ OLs and Sox10^+^CC1^-^ OPCs (Fig. 3J) in the spinal cord of P9 *Pdgfra:Parp1* cKO and Ctrl mice which had been administered with tamoxifen at P1, P2, and P3. The density of OLs was significantly diminished in *Pdgfra:PARP1* cKO spinal cord, which was corroborated by reduced mRNA levels of *Mbp* and *Qk* (antigen recognized by clone CC1) (Fig. 3K). Consistent with the defective OPC differentiation, the expression of newly formed OL marker genes (Fig. 3L) was significantly lower in the spinal cord of *Pdgfra:Parp1* cKO mice than that of *Parp1* controls, suggesting a diminished oligodendrogenesis rate. Thus, these data conclusively demonstrate that PARP1 is required for OPC differentiation in the murine CNS.

### Oligodendroglial PARP1 is essential for developmental myelination

To determine the necessity of PARP1 in CNS myelination, we analyzed oligodendroglial myelin formation in the corticospinal tract (CST) of P14 *Pdgfrα:Parp1* cKO mice (Fig. 3M). Analysis for the light microscopic images of toluidine blue myelin staining demonstrated a significant decrease of myelinated axon number in *Pdgfrα:Parp1* cKO mice (Fig. 3N). Ultra-structural myelin assays for TEM images demonstrated that the number of myelinated axons (diameter ≥ 0.3μm) was significantly decreased in *Pdgfrα:Parp1* cKO mice (Fig. 3O), corroborating toluidine blue myelin staining assay. The density of total axons (myelinated + unmyelinated) with diameter ≥ 0.3μm was comparable between *Pdgfrα:Parp1* cKO mice and controls (Fig. 3P). Among the myelinated axons, the myelin sheath was significantly thinner, as evidenced by greater g-ratio in *Pdgfrα:Parp1* cKO mice than that in *Parp1* controls (Fig. 3Q-R). Furthermore, the caliber distribution of myelinated axons appeared comparable between *Pdgfrα:Parp1* cKO and Ctrl mice (data now shown), suggesting that *Pdgfrα:Parp1* cKO does not affect axonal physical properties. These data collectively demonstrate that PARP1 is required for oligodendroglial myelination.

### PARP1 is activated primarily in OLCs during postnatal CNS development

To elucidate the mechanisms underlying PARP1-regulated oligodendroglial development, we sought to interrogate if PARP1’s activity plays a part. It remains to be defined whether PARP1 is activated in neural cells during postnatal CNS development. To this end, we employed PAR immune-detection as a reliable surrogate for PARP1’s enzymatic activation (Fig. 4A) (Luo and Kraus, 2012). We found that the nuclear PAR immunoreactive signals were present in the spinal cord of P7 mice (Fig. 4B, arrowheads) and absent in the adult CNS (data not shown, cf Fig. 7B). Immunoreactive PAR signals were abolished in the spinal cord of age-matched *Parp1* KO mice, as evidenced by IHC (Fig. 4B, bottom). These data suggest that PARP1 is the major PARP family member responsible for cellular PARylation activity in the CNS and that PARP1’s enzymatic activity is dynamically regulated during postnatal CNS development.

**Fig. 4.**
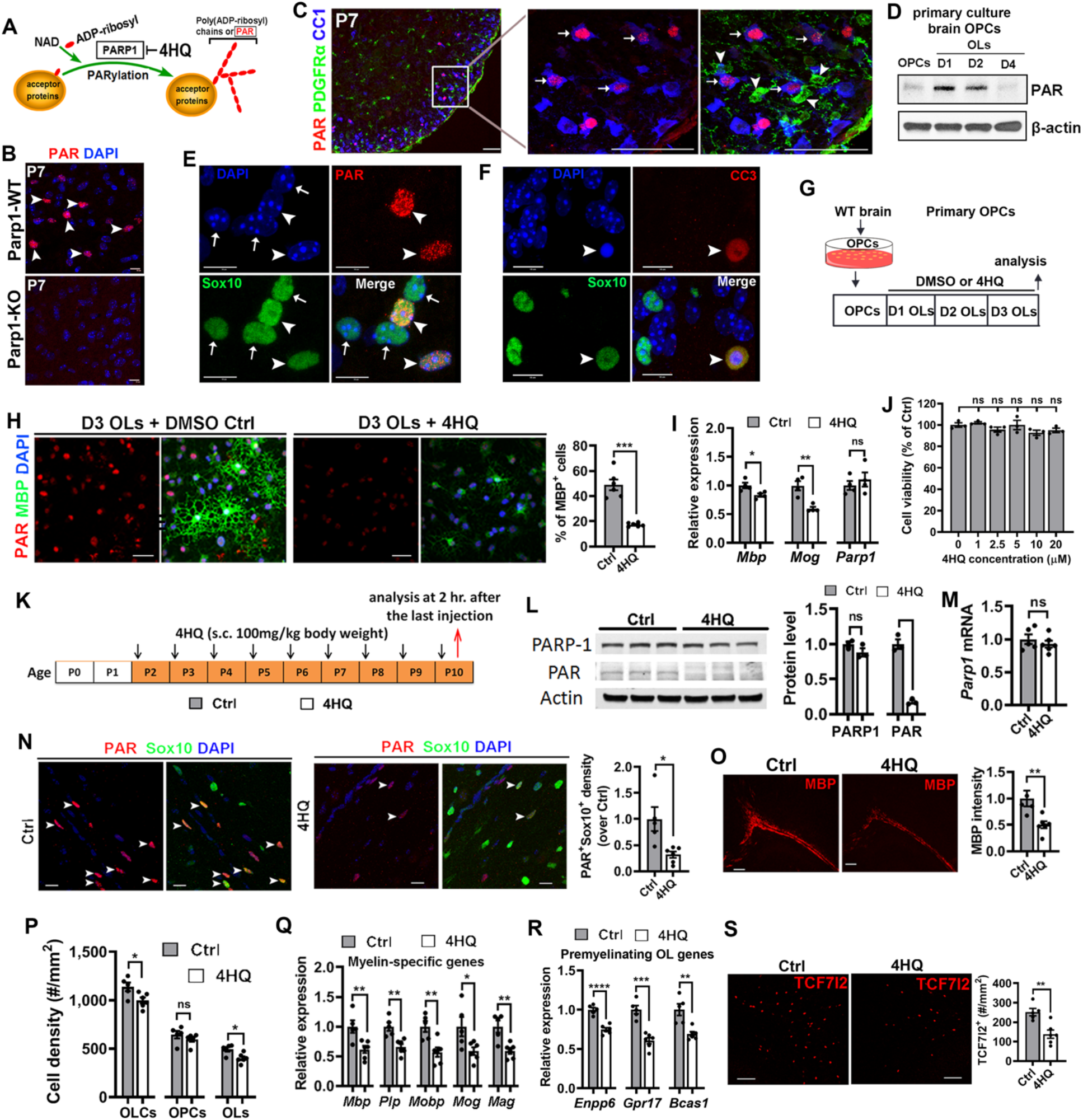
PARP1 activity is essential for OPC differentiation *in vitro* and *in vivo*. **A,** upon enzymatic activation, PARP1 catalyzes the covalent addition of poly (ADP-ribosyl) units (PAR) to acceptor proteins using NAD^+^ molecules as PAR donor. Thus, PAR immune-detection is reliable function readout of PARP1’s enzymatic activity. 4- hydoxyquinazoline (4HQ) inhibits PARP1 activity, leading decreased PAR deposits on acceptor proteins. **B**, immunohistochemical detection of PAR in P7 spinal cord. Arrowheads, nuclear PAR-positive cells. Note that PAR signals were undetectable in Parp1 KO mice. **C**, triple IHC of PAR, PDGFRa, and CC1 in P7 spinal cord. Arrowheads, PDGFRa^+^ OPCs; arrows, CC1^+^PAR^+^ OLs. **D**, Western blot assay for PAR at ∼116kD using protein extracts from primary brain OPCs and differentiating OLs at D1, D2, and D4. **E**, high-power confocal images showing the nuclear morphology (indicated by DAPI staining) of PAR^+^Sox10^+^ (arrowheads) and PAR^-^Sox10^+^ (arrows) oligodendrocytes in P10 spinal white matter. **F**, IHC of Sox10 and active cleaved caspase 3 (CC3) showing the condensation of DAPI^+^ nucleus of an apoptotic oligodendrocyte (arrowheads) in P10 spinal white matter. **G**, experimental design for panel **H-J**. 4HQ (10 µM) was included in the differentiation medium for 3 days. **H**, confocal images showing decreased PAR immunoreactive signals and increased MBP expression, and percent of MBP^+^ OLs with ramified morphology among DAPI^+^ nuclei. N = 6 Ctrl, 6 4HQ. Welch-corrected *t*_(5.129)_ = 7.362 *P* = 0.0006. **I**, qRT-PCR assay for *Mbp*, *Mog, and Parp1*. N = 4 Ctrl, 4 4HQ. *Mbp*, *t*_(6)_ = 2.981 *P* = 0.0246; *Mog*, *t*_(6)_ = 4.582 *P* = 0.0038; *Parp1*, *t*_(6)_ = 0.7001 *P* = 0.5101. **J**, MTT assay for cell viability upon 4HQ treatment in differentiation medium at different doses up to 20 µM. One-way ANOVA followed by Tukey’s multiple comparison test. *F*_(5,12)_ = 1.977, *P* = 0.1548, ns, not significant. **K**, experimental designs for Panel **L-S**. Data were collected from the brain. **L**, Western blotting and quantification of PARP1 and PAR (at ∼116kD). N = 3 Ctrl, 3 4HQ. PARP1, *t*_(4)_ = 1.734 *P* = 0.1580; PAR, *t*_(4)_ = 11.37 *P* = 0.0.0003. **M,** *Parp1* mRNA in the brain quantified by qRT-PCR. N = 5 Ctrl, 6 4HQ. *t*_(9)_ = 0.832 *P* = 0.4269. **N,** representative confocal images and quantification of PAR and Sox10 in the corpus callosum. Arrowheads, PAR^+^Sox10^+^ oligodendroglial cells. N = 5 Ctrl, 6 4HQ. Welch-corrected *t*_(4.556)_ = 2.875 *P* = 0.0388. **O**, myelination visualized by MBP staining in the corpus callosum and external capsule (left) and relative MBP intensity (right). N = 5 Ctrl, 6 4HQ. *t*_(9)_ = 3.287 *P* = 0.0094. **P**, density of Sox10^+^ OLCs, Sox10^+^PDGFRα^+^ OPCs, and Sox10^+^CC1^+^ OLs in the corpus callosum. N = 5 Ctrl, 6 4HQ. OLCs, *t*_(9)_ = 2.656 *P* = 0.0262; OPCs, *t*_(9)_ = 1.131 *P* = 0.2873; OLs, *t*_(9)_ = 3.027 *P* = 0.0143. **Q**, qRT-PCR assays for major myelin-specific genes. N = 5 Ctrl, 6 4HQ. *Mbp*, *t*_(9)_ = 3.300 *P* = 0.0092; *Plp*, *t*_(9)_ = 4.073 *P* = 0.0028; *Mobp*, *t*_(9)_ = 3.576 *P* = 0.0060; *Mog*, *t*_(9)_ = 2.536 *P* = 0.0319; *Mag*, *t*_(9)_ = 3.841 *P* = 0.0040. **R**, qRT-PCR assays for genes enriched in premyelinating OLs. N = 5 Ctrl, 6 4HQ. *Enpp6*, *t*_(9)_ = 7.092 *P* < 0.0001; *Gpr17*, *t*_(9)_ = 6.153 *P* = 0.0002; *Bcas1*, *t*_(9)_ = 4.147 *P* = 0.0025. **S**, representative confocal images of TCF7l2 (left) and density of TCF7l2-expressing newly differentiated OLs (right). N = 5 Ctrl, 6 4HQ. *t*_(9)_ = 3.897 *P* = 0.0036. Unpaired, two-tailed Student’s *t* test, \**p* < 0.05; \*\**p* < 0.01; \*\*\**p* < 0.001; ns, not significant *p* > 0.05. Scale bars: B, E, F, 10μm; H, N, 20µm; C, 50μm; O, 200µm.

We next used lineage-specific markers to determine what cell type exhibits PARP1 activity. Double IHC demonstrated that the vast majority of PAR^+^ cells (94.5 ± 5.2%, n = 3) were Sox10-expressing oligodendroglial lineage cells in the spinal white matter at P7. Triple IHC showed that PAR signals were low in PDGFRα^+^ OPCs (Fig. 4C, arrowheads), upregulated in CC1^+^ OLs (Fig. 4C, arrows) in the spinal cord at P7, and undetectable in CC1^+^ OLs (data not shown) in the adult spinal cord. Using primary oligodendrocyte cultures purified from rodent brain and Western blotting assay, we demonstrated that PAR signals were at the barely detectable level in OPCs, transiently upregulated in newly formed immature OLs (D1 and D2) and downregulated in mature OLs (D4) (Fig. 4D). Thus, PARP1 is transiently activated primarily in oligodendroglial lineage cells during postnatal CNS development.

Sustained activation of PARP1 in response to excessive DNA damaging agents leads to cell apoptosis in a context-dependent manner (Alano et al., 2010; Hassa, 2009; Scott et al., 2003). We found that the nuclear morphology of PAR^+^Sox10^+^ cells (Fig. 4E, arrowheads) was indistinguishable from that of PAR^-^Sox10^+^ cells (Fig. 4E, arrows), both of which exhibited no morphological characteristics of dying cells, such as nuclear condensation (Fig. 4F, arrowheads). These data suggest that PARP1 activation in oligodendroglial lineage cells may play other roles than cell death in the developing CNS with minimal or no genotoxic injury.

### PARP1 enzymatic activity is required for OPC differentiation

In the CNS of *Parp1* KO (Fig. 2) and *Parp1* cKO (Fig. 3) mice, both activity-dependent and independent function of PARP1 is eliminated. To elucidate the function of PARP1 activity in oligodendroglial development, we used the potent PARP1 inhibitor, 4-hydoxyquinazoline (4HQ), to dampen cellular PARylation (Fig. 4A) and determine the cell-autonomous effect of PARP1 inhibition on OPC differentiation in purified primary OPC cultures (Fig. 4G). We demonstrated that 4HQ treatment remarkably reduced PAR signals (Fig. 4H) without affecting PARP1 expression (Fig. 4I). The percent of ramified MBP^+^ OLs was significantly decreased in 4HQ-treated group (Fig. 4H), indicating an inhibition of OPC differentiation and maturation. Consistently, the expression of myelin-specific genes was significantly lower in 4HQ-treated group than that in Ctrl group (Fig. 4I). MTT cell viability assay demonstrated that the 4HQ concentration we used (10 μM) had no effect on cell death (Fig. 4J). These *in vitro* data demonstrate that the defective OPC differentiation in 4HQ-treated OPC culture phenocopies that in PARP1-deficient OPC culture (Fig. 2O-Q), which suggests that PARP1 activity-dependent function plays a crucial role in PARP1-regulated OPC differentiation.

We demonstrated that PARP1 was activated predominantly in OLCs *in vivo*. We next determined whether PARP1 activity plays a role OPC differentiation *in vivo*. 4HQ administration to neonatal mice (Fig. 4K) significantly decreased the number of Sox10^+^ OLCs that are positive for PAR in the corpus callosum (Fig. 4L, N) without perturbing PARP1 expression (Fig. 4L, M). Myelination, visualized by MBP histological staining, was significantly reduced in the subcortical white matter of 4HQ-treated mice (Fig. 4O). Quantification data showed that the density of CC1^+^ differentiated OLs was significantly decreased in 4HQ-treated mice (Fig. 4P) and that the expression of myelin-specific genes was much lower in the brain of 4HQ-treated mice than that in controls (Fig. 4Q). Moreover, the expression of pre-myelinating OL-enriched genes (Fig. 4R) and the density of TCF7l2^+^ premyelinating OLs (Fig. 4S) were markedly diminished in the brain of 4HQ-treated mice, suggesting that PARP1 inhibition diminishes brain oligodendrogenesis rate *in vivo*. We further showed that the number of OLs was significantly diminished in the spinal cord of 4HQ-treated mice compared with that in control spinal cord (fig. S5). Western blotting assay revealed no evidence for alterations of cell death including PARP1-mediated parthanatos, Gasdermin D-mediated pyroptosis, and phosphorylated MLKL-mediated necrosis (Galluzzi et al., 2012) in 4HQ-treated brains (fig. S6A-B). We found no evidence of microglial (fig. S6C-E) or astroglial (fig. S6F-H) activation in the brain of 4HQ-treated mice compared with age-matched control mice at the histological and molecular levels, which is consistent with minimal or no PARP1 activity in microglial and astroglial cells. Collectively, these data suggest that PARP1 enzymatic activity is essential for OPC differentiation in a cell autonomous manner and that PARP1 activation plays a minor role in mediating cell death and glial activation during developmental myelination in the CNS.

### Stabilizing PARylation is sufficient to promote OPC differentiation

The half-life of PAR chains deposited on acceptor proteins by PARP1 is less than 1 minute (Zhen and Yu, 2018). The rapid PAR turnover is catalyzed by the PAR degrading enzyme PARG (Fig. 5A). Having shown that disrupting PARP1 and inhibiting PARP1-mediated PARylation impede oligodendroglial development, we next asked if stabilizing PARylation (PARP1 gain-of-function) affects OPC differentiation during postnatal CNS development. To this end, we treated neonatal mice with a potent PARG inhibitor, PDD 00017273 (PDD, thereafter) (Gogola et al., 2018; James et al., 2016) (Fig. 5B) to augment PARylation through dampening PAR degradation (Fig. 5C). PDD treatment increased the number and intensity of PAR immunoreactive signals in Sox10^+^ OLCs in the spinal cord and forebrain corpus callosum (Fig. 5D). Strikingly, we found that the density of CC1^+^ OLs was significantly increased in the ventral white matter of spinal cord and the forebrain corpus callosum of PDD-treated mice compared with that of control mice (Fig. 5E), whereas PDGFRα^+^ OPCs were not affected (Fig. 5F). These data indicate that augmenting cellular PARylation level promotes oligodendrogenesis.

**Fig. 5.**
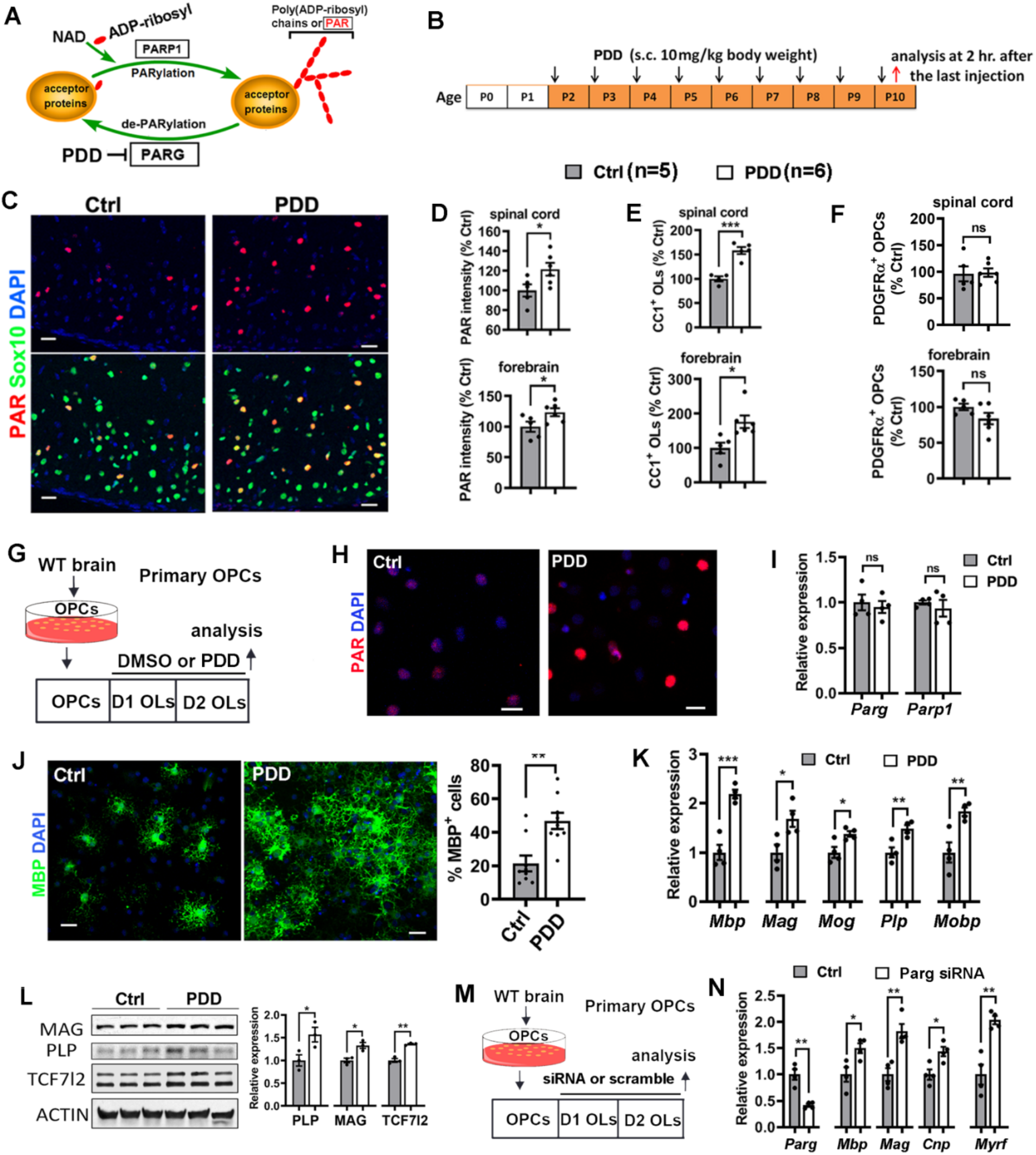
Stabilizing cellular PARylation promotes OPC differentiation *in vivo and in vitro.* **A**, diagram depicting the stabilization of cellular PARylation by inhibiting the PAR degrading enzyme PARG. **B**, experimental designs for Panel **C-F**. **C**, representative confocal images of PAR and Sox10 in the spinal cord ventral white matter of PDD treated and vehicle control mice. **D**, quantification of relative PAR intensity in the spinal cord white matter (upper) and forebrain corpus callosum (lower). Spinal cord *t*_(9)_ = 2.357 *P* = 0.0428; forebrain *t*_(9)_ = 2.291 *P* = 0.0477. **E**, relative density of CC1^+^ OLs in the spinal cord white matter (upper) and forebrain corpus callosum (lower). Spinal cord *t*_(9)_ = 6.209 *P* = 0.0004; forebrain *t*_(9)_ = 3.101 *P* = 0.0127. **F**, relative density of PDGFRα^+^ OPCs in the spinal cord white matter (upper) and forebrain corpus callosum (lower). Spinal cord *t*_(9)_ = 0.1519 *P* = 0.8826; forebrain *t*_(9)_ = 1.664 *P* = 0.1304. **G**, experimental designs for Panel **H-L**. PDD concentration in the differentiation medium was 1 μM. **H**, confocal images of PAR showing increased intensity and number of PAR-positive cells in PDD-treated group. **I**, qRT-PCR assay showing comparable level of *Parg* and *Parp1*. N = 4 Ctrl, 4 PDD. *Parg*, *t*_(6)_ = 0.4556 *P* = 0.6647; *Parp1*, *t*_(6)_ = 0.6733 *P* = 0.5258. **J**, representative ICC images (left) and percent of ramified MBP^+^ OLs among DAPI^+^ cells (right). N = 9 Ctrl, 9 PDD. *t*_(16)_ = 3.736 *P* = 0.0018. **K**, mRNA levels of myelin genes quantified by qRT-PCR. N = 4 Ctrl, 4 PDD. *Mbp*, *t*_(6)_ = 6.691 *P* = 0.0005; *Mag*, *t*_(6)_ = 2.994 *P* = 0.0242; *Mog*, *t*_(6)_ = 2.984 *P* = 0.0245; *Plp*, *t*_(6)_ = 3.822 *P* = 0.0087; *Mobp*, *t*_(6)_ = 3.719 *P* = 0.0099. **L**, Western blotting images (left) and quantification (right) of myelin proteins MAG, PLP, and pre-myelinating OL marker TCF7l2. β-actin, loading control. N = 3 Ctrl, 3 PDD. PLP, *t*_(4)_ = 2.805 *P* = 0.0485; MAG, *t*_(4)_ = 4.050 *P* = 0.0155; TCF7l2, *t*_(4)_ = 8.176 *P* = 0.0012. **M**, primary OPCs purified from neonatal rat brain were differentiated in the presence of *Parg* siRNA or scramble control for 2 days prior to analysis. **N**, relative expression of *Parg*, myelin-specific genes and the potent pro-differentiation factor *Myrf* quantified by qRT-PCR. N = 4 Ctrl, 4 *Parg* siRNA. *Parg, t*_(6)_ = 5.650 *P* = 0.0013; *Mbp*, *t*_(6)_ = 2.995 *P* = 0.0242; *Mag*, *t*_(6)_ = 4.447 *P* = 0.0043; *Cnp*, *t*_(6)_ = 3.312 *P* = 0.0163; *Myrf*, *t*_(6)_ = 4.990 *P* = 0.0025. Two-tailed Student’s t test, \**P* < 0.05; \*\**P* < 0.01; \*\*\**P* < 0.001; ns, not significant. Scale bars: C, 25 μm; H, J, 10 μm.

To determine whether stabilizing PARylation affects OPC differentiation in a cell-autonomous manner, we sought to employ purified primary OPC culture (Fig. 5G). PDD treatment markedly increased PAR immunoreactive signals (Fig. 5H) without altering the expression of PARP1 and PARG (Fig. 5I). MTT assay showed that PDD did not affect the cell death or survival of primary oligodendrocytes at the dose we used (data not shown). We found that PDD inhibition markedly increases the percent of MBP-expressing OLs bearing arborized morphology compared with Ctrl group at 36 hours post-treatment (Fig. 5J). To corroborate this observation, qRT-PCR assay showed that the mRNA levels of major myelin proteins were significantly higher in PDD group than those in Ctrl group (Fig. 5K). Western blotting assay demonstrated that the protein level of myelin protein MAG and PLP and newly formed OL marker TCF7l2 was significantly increased in PDD-treated group (Fig. 5L), suggesting that inhibiting PARG activity promotes OPC differentiation and myelin gene expression. To strengthen the conclusion, siRNA-mediated PARG knockdown in purified primary rat OPCs (Fig. 5M) significantly increased the expression of major myelin genes (*Mbp, Cnp, and Mag*) and the potent pro-differentiation factor *Myrf* (Fig. 5N), suggesting that PARG (and its activity) regulates OPC differentiation in a cell-autonomous manner. Taken together, our data demonstrate that stabilizing PARP1’s PARylation function by inhibiting PARG activity or decreasing PARG expression is sufficient to promote OPC differentiation.

### Unbiased proteomic analysis identifies novel target proteins of PARP1 activity

PARP1 activity regulates cellular processes mainly through binding and modulating its PARylated target proteins (Luo and Kraus, 2012). Having demonstrated that PARP1-mediated PARylation is necessary and sufficient for OPC differentiation and OL maturation, we sought to identify potential PAR acceptor (target) proteins, providing further molecular mechanisms underlying PARP1-regulated oligodendroglial development. To this end, PAR antibody-mediated Co-IP was used to enrich PARylated proteins from P7 spinal cord where PAR^+^ signals are observed selectively in oligodendrocytes (Fig. 4C), followed by protein identification by LC-MS/MS (Fig. 6A). We screened 131 potential candidates of PARylated nuclear proteins that were significantly enriched by at least 3-fold (P < 0.05) in PAR antibody-IP components compared with IgG Ctrl-IP components. GO analysis for biological processes (GO_BP) and molecular function (GO_MF) showed that the identified PAR acceptor proteins were predominantly associated with mRNA processing, splicing, transporting, and translation (Fig. 6B) and participated in RNA binding function (Fig. 6C). This is in stark contrast to the PARylated proteins previously identified from cells treated by the genotoxic agent H_2_O_2_ (Zhang et al., 2013) which were primarily related to DNA damage and repair (fig. S7A-B). These data suggest that PARP1’s enzymatic activity may play a minor role, if any, in genotoxin-induced cell death under physiological conditions in the CNS where genome stability is maintained.

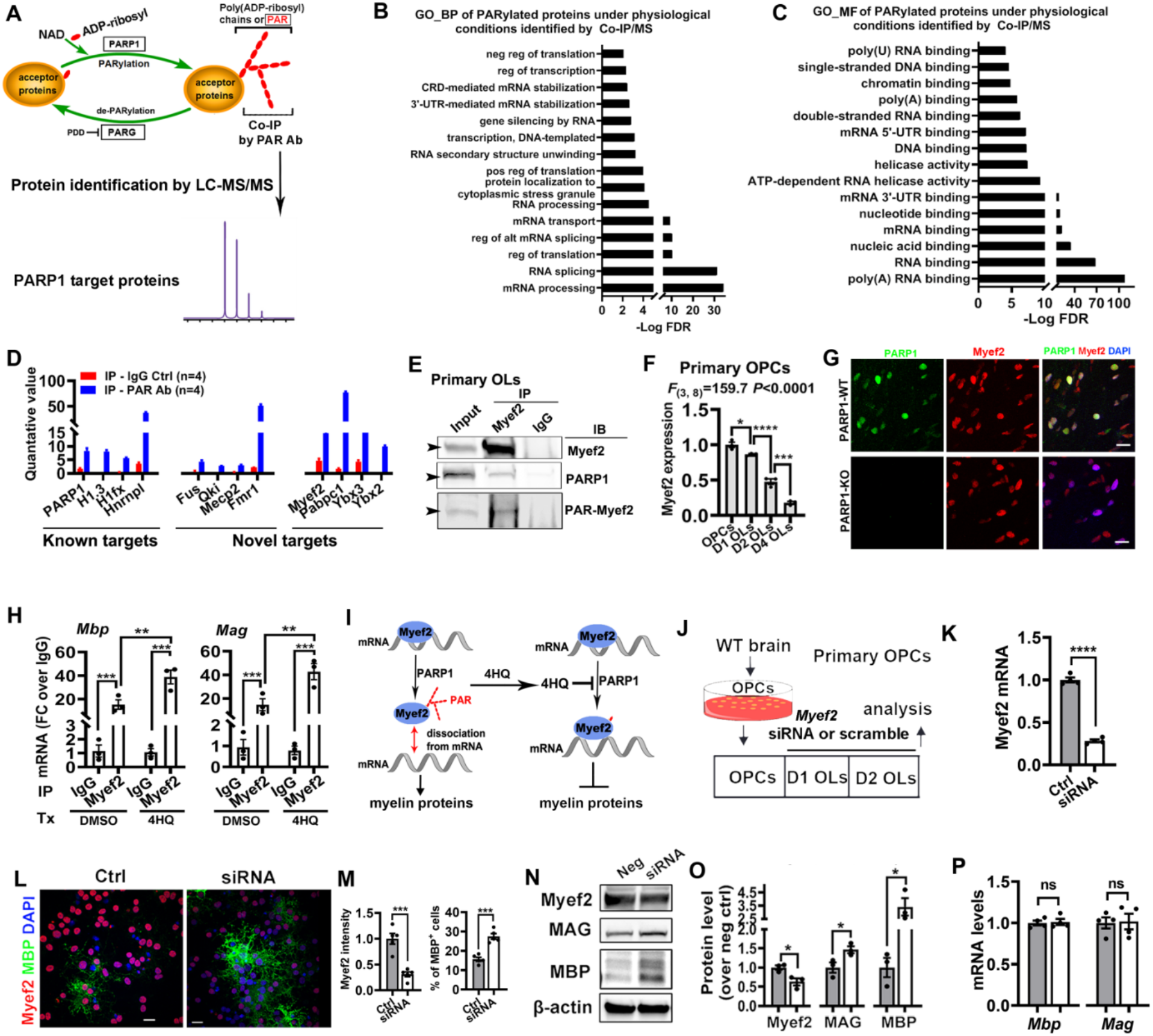
PARP1 PARylates Myef2 to regulate OPC differentiation and myelin gene expression. **A,** flow chart of identifying PARylated acceptor proteins. Whole proteins were prepared from the spinal cord of P7 mice 30 min after PARG inhibitor PDD injection to stabilize protein PARylation (PAR half-life is less than 1min). **B-C**, GO analysis of biological processes (**B**) and molecular function (**C**) of the 131 potential PARylated acceptor proteins identified by Co-IP and LC-MS/MS. **D,** representative graphs depicting examples of known PARylated proteins and novel PARylated proteins. **E,** Co-IP followed by Western blotting assay confirming the interaction between Myef2 and PARP1 and the presence of PARylated Myef2 at the molecular weight of Myef2 (∼55kD) in the primary brain OLs at D2. Rabbit IgG, negative control for Myef2 IP. **F,** Myef2 mRNA level in primary OPCs and D1, 2 and 4 differentiating OLs. N = 3 each group. One-way ANOVA followed by Tukey’s multiple comparison tests. **G,** double IHC of PARP1 and Myef2 in the corpus callosum of P7 Parp1-WT and Parp1-KO mice. **H**, abundance of *Mbp* and *Mag* mRNA in the immunoprecipitates using RNA immunoprecipitation (RIP) by Myef2 antibody or Ctrl IgG in D2 primary OLs treated with DMSO or 4HQ treated. N = 3 each group. Two-way ANOVA followed by Sidak multiple comparison tests. *Mbp:* DMSO vs 4HQ *F*_(1, 8)_ = 10.79 *P* = 0.0111, IgG vs Myef2 antibody *F*_(1, 8)_ = 53.51 *P* < 0.0001. *Mag:* DMSO vs 4HQ *F*_(1, 8)_ = 10.82 *P* = 0.0110, IgG vs Myef2 antibody *F*_(1, 8)_ = 43.75 *P* = 0.0002. **I**, schematic conclusion of panel **H**, PARP1 inhibition by 4HQ potentiated the binding of Myef2 to mRNA molecules. **J**, experimental designs for Panels **K-P**. Primary OPCs purified from neonatal rat brain were differentiated in the presence of *Myef2* siRNA or scramble controls for 36 h in the differentiation medium prior to analysis. **K,** qRT-PCR assay for *Myef2* mRNA level. N = 4 Ctrl, 4 *Myef2* siRNA. *Myef2, t*_(6)_ = 21.36 *P* <0.0001. **L**, ICC of Myef2 and MBP in Ctrl and *Myef2* siRNA-treated primary OLs. **M**, quantification of Myef2 intensity and MBP^+^ ramified OLs (percentage of DAPI^+^ cells). N = 5 Ctrl, 4 *Myef2* siRNA. Myef2 intensity*, t*_(8)_ = 6.084 *P* = 0.0003; % of MBP^+^ cells, *t*_(8)_ = 6.672 *P* = 0.0002. **N-O,** representative Western blotting images (**N**) and quantification (**O**). β-actin, loading control. N = 3 scramble Ctrl, 4 *Myef2* siRNA. Myef2*, t*_(4)_ = 3.410 *P* = 0.0270; MAG, *t*_(4)_ = 2.893 *P* = 0.0444; MBP, *t*_(4)_ = 3.387 *P* = 0.0276. **P,** qRT-PCR assay for *Mbp* and *Mag* in Ctrl (n = 4) and *Myef2* siRNA-treated (n = 4) primary OLs. *Mbp, t*_(6)_ = 0.2572 *P* = 0.8056; *Mag*, *t*_(6)_ = 0.1497 *P* = 0.8859. Two-tailed Student’s t test, \**P* < 0.05; \*\**P* < 0.01; \*\*\**P* < 0.001; ns, not significant. Scale bars: G, 20 μm; L 10 μm.

Amongst the PARylated protein candidates were well-known acceptor proteins including PARP1, the primary PAR acceptor protein in cells (Luo and Kraus, 2012), histone protein (Histone H1.3 and H1fx), and heterogeneous nuclear ribonucleoproteins (hnRNPs) (Ke et al., 2019) (Fig. 6D). The identification of these proteins provided positive controls for our unbiased approaches. Very interestingly, we identified some novel PARylated proteins with reported functions in oligodendroglial development, such as the RNA/DNA-binding protein Fus (Guzman et al., 2020) and Mecp2 (Nguyen et al., 2013) and the RNA-binding protein Qki (Thangaraj et al., 2017; Zhou et al., 2020) and Fmr1 (Giampetruzzi et al., 2013; Shi et al., 2019) (Fig. 6D). We also identified novel PARP1 target proteins with yet unknown functions in oligodendroglial development, such as the myelin expression factor 2 (Myef2), Y-box binding proteins (Ybx3, Ybx2), and polyadenylate-binding protein 1 (Pabpc1) (Fig. 6D) among others. Together, our data suggest a potential role of PARP1 in regulating RNA metabolism and RNA binding under physiological conditions and provide valuable novel PARP1 target proteins for future studies.

### Myef2 is a novel PARP1 target, negatively regulating oligodendroglial differentiation

To further validate our unbiased proteomic identification, Co-IP followed by Western blotting assay confirmed that Myef2 interacted with PARP1 and was PARylated by PARP1 in primary OLs (Fig. 6E). Using rodent primary culture, we showed that Myef2 expression was downregulated along oligodendroglial lineage progression and maturation (Fig. 6F). We found that genetic *Parp1* KO did not affect Myef2 expression at the protein (Fig. 6G) and mRNA (data not shown) levels. These data suggest that, instead of regulating its expression, PARP1 may modulate Myef2’s function through adding bulky negatively charged PAR chains.

Recent data suggest that Myef2 binds to the majority mRNAs of protein-coding genes in oligodendroglial cell lines through 3’-untranslated region (UTR) (Samudyata et al., 2019) which often involves in repressing mRNA processing and translation (Abaza and Gebauer, 2008; Szostak and Gebauer, 2013). We hypothesize that Myef2 may bind myelin gene mRNAs and regulate myelin gene expression at the post-transcriptional level. We employed RNA immunoprecipitation approach (He et al., 2017) to pull down Myef2-bound mRNA from D2 primary OLs. Our data demonstrated that myelin gene mRNA transcripts of *Mbp* and *Mag* were markedly enriched in Myef2 antibody components compared to IgG control components (Fig. 6H), suggesting that Myef2 binds to myelin gene transcripts. Notably, inhibiting PARP1’s PARylation activity by 4HQ significantly enhanced the binding of Myef2 to MBP and MAG mRNAs (Myef2+DMSO vs Myef2+4HQ) (Fig. 6H). These data indicate that PARP1-mediated Myef2 PARylation dissociates Myef2 from binding myelin gene transcripts in oligodendroglial lineage cells (Fig. 6I).

The function of Myef2 in myelin gene expression remains undefined. To determine its role in oligodendroglial development, we knocked down Myef2 in rodent primary OPC cultures by siRNA (Fig. 6J-K). We found that the percent of MBP^+^ cells with ramified morphology (Fig. 6L) was increased by ∼80% in *Myef2* siRNA cultures compared with scrambled controls (Fig. 6M). Consistently, Western blotting assay (Fig. 6N) demonstrated that myelin gene protein MBP and MAG were significantly increased in *Myef2* siRNA cultures (Fig. 6O). In sharp contrast, the mRNA levels of *Mbp* and *Mag* were not affected by *Myef2* knockdown (Fig. 6P), suggesting that Myef2 negatively regulates OPC differentiation by repressing myelin gene expression at the post-transcriptional level. Together, these data indicate that PARP1’s PARylation activity positively regulates OL differentiation at least in part through PARylating Myef2 and subsequently relieving the repressive effect of Myef2 on myelin gene expression at post-transcriptional level (Fig. 6I).

### PARP1 activity is induced in cuprizone-induced demyelination and remyelination

To determine the significance of PARP1 activity in OL regeneration and remyelination, we employed the mouse model of cuprizone (CPZ)-induced demyelination (Fig. 7A) in which 6 weeks of CPZ diet results in complete demyelination in the corpus callosum (CC) followed by full remyelination by 6 weeks after animal returning to the normal diet (Kipp et al., 2009; Matsushima and Morell, 2001). We observed that PARP1 expression was barely detectable except for low level in PDGFRα^+^ OPCs (Fig. 7B, left) and that PAR was absent (Fig. 7B, right) from the CC of adult mice maintained on the normal diet. However, during remyelination process, PARP1 was markedly induced in CC1^+^ newly regenerated OLs (Fig. 7C). Similarly, PARP1’s PARylation activity, evidenced by PAR IHC, was elevated predominantly in CC1^+^ newly regenerated OLs at 6+1 wk (Fig. 7D) and downregulated to the barely detectable level at 6+6 weeks (Fig. 7E) when full remyelination has been achieved (Kipp et al., 2009; Matsushima and Morell, 2001). The temporal dynamics of PARP1 activity during remyelination is reminiscent of that during developmental myelination (Fig. 4) and suggests that PARP1 activity may regulate OL regeneration and myelin repair.

**Fig. 7.**
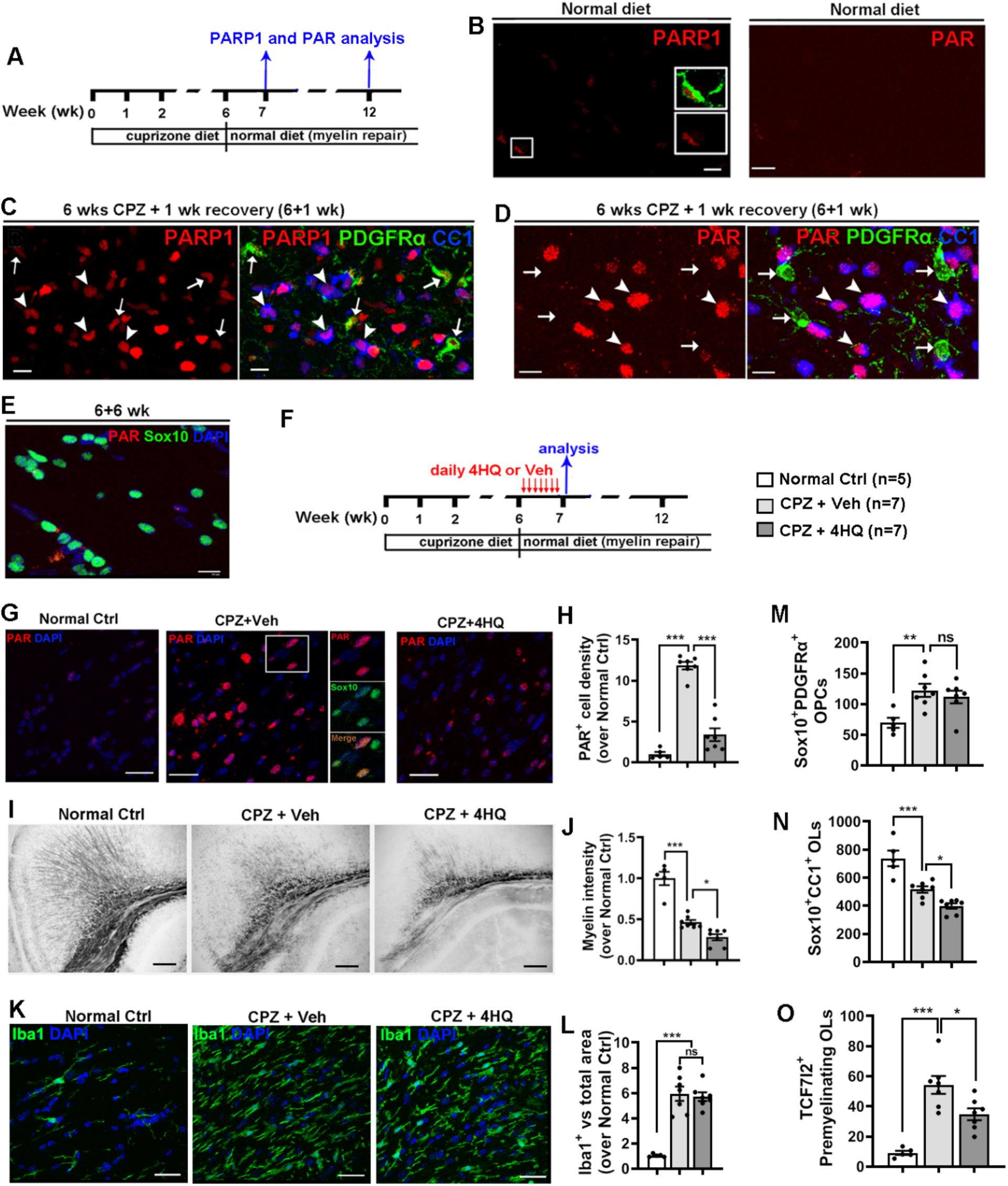
PARP1 activity is required for oligodendrocyte regeneration and remyelination. **A,** experimental design for panel **B-E**. 2-month old C57BL/6J mice were fed with 0.2% Cuprizone (CPZ) diet for 6 weeks for complete demyelination followed by 6 week with normal diet for complete remyelination. **B**, IHC showing low level of PARP1 expression and absence of PAR signals in the adult corpus callosum (CC) of mice maintained on normal diet. Boxed area was shown at higher magnification, green, PDGFRα. **C**, IHC of PARP1/CC1/PDGFRα in the CC of CPZ-treated mice at 1 week after returning to the normal diet (6+1 wk). Arrows, PARP^+^PDGFRα^+^ OPCs; arrowheads, PARP^+^CC1^+^ newly regenerated OLs. **D**, IHC of PAR/CC1/PDGFRα in the CC at 6+1 wk. Arrows, PARP^-^PDGFRα^+^ OPCs; arrowheads, PAR^+^CC1^+^ newly regenerated OLs. **E**, IHC of PAR/Sox10 in the CC at 6+6 wk. **F,** experimental design for panel **G-O**. 4HQ (100 mg/kg) and DMSO vehicle (Veh) were i.p. injected daily starting from day 1 through day 7 after returning to the normal diet, and the posterior corpus callosum was processed for analysis. **G-H**, representative IHC images (**G**) and quantification (**H**) of PAR^+^Sox10^+^ cells in the CC. Boxed area was shown for Sox10 and PAR individual channels. **I-J**, black gold myelin staining (**I**) and quantification (**J**) of remyelination in the CC and overlaying cortex. **K-L**, representative confocal images of Iba1^+^ activated microglia/macrophages (**K**) and quantification (**L**) in the CC. **M-O**, quantification (#/mm^2^) of Sox10^+^PDGFRα^+^ OPCs, Sox10^+^CC1^+^ OLs, and TCF7l2^+^ premyelinating OLs in the CC. Statistics: one-way ANOVA with Tukey’s multiple comparison test, \**P* < 0.05, \*\**P* < 0.01, \*\*\**P* < 0.001, ns, not significant. **H**, *F*_(2, 16)_ = 90.75, *P* < 0.0001; **J**, *F*_(2, 16)_ = 55.37, *P* < 0.0001; **L**, *F*_(2, 16)_ = 36.98, *P* < 0.0001; **M**, *F*_(2, 16)_ = 6.615, *P* = 0.0081; **N**, *F*_(2, 16)_ = 26.91, *P* < 0.0001; **O**, *F*_(2, 16)_ = 22.01, *P* < 0.0001. Scale bars, B-E, 10µm; G, K, 20µm, I, 50µm.

### Inhibiting PARP1’s activity impairs oligodendrocyte regeneration and remyelination

To define the role of PARP1 enzymatic activity in remyelination, we treated CPZ-demyelinated mice with 4HQ or vehicle DMSO (Veh) on day 1 after returning to the normal diet for 7 consecutive days, and the treated mice were euthanized 2 h after the last 4HQ injection (Fig. 7F), a time point that active oligodendrogenesis and remyelination occur in the CC (Hammond et al., 2015). 4HQ treatment significantly reduced PARP1’s PARylation activity (Fig. 7G), as evidenced by significant diminution in immunoreactive PAR signals (Fig. 7H). Black gold II myelin staining (Fig. 7I) demonstrated that PARP1 inhibition significantly impaired remyelination in the CC of CPZ+4HQ group compared with that of CPZ+Veh group (Fig. 7J). We found that the 4HQ-treatment paradigm did not affect the activation of microglia/macrophages quantified by Iba1 (Fig. 7K) compared with that in CPZ+Veh group, both of which were upregulated by ∼ 6-fold (Fig. 7L) in comparison with normal Ctrl group, suggesting that the myelin debris clearance function of activated microglia/macrophages (Cignarella et al., 2020; Lampron et al., 2015) is not affected by 4HQ.

Defective myelin repair implies that the extent of oligodendrocyte regeneration from activated OPCs is impaired in 4HQ-treated group. We next used developmental stage-dependent markers to quantify different maturation stages of OLCs in the CC. Our quantification showed that the number of Sox10^+^PDGFRα^+^ OPCs was similar between 4HQ and Veh-treated groups, both of which were significantly higher than that in normal Ctrl mice (Fig. 7M), suggesting that PARP1 activity is dispensable for OPC activation and population expansion. In contrast, the number of Sox10^+^CC1^+^ total OLs was significantly decreased in the CC of 4HQ group compared with that of Veh group (Fig. 7N). Consistently, the number of TCF7l2^+^ newly regenerated premyelinating OLs at the time of mouse sacrifice was diminished in group compared with that of Veh group (Fig. 7O), indicating that PARP1 inhibition decrease the rate of oligodendrogenesis from OPCs. Together, our data suggest that PARP1 activity is required for oligodendrocyte regeneration and remyelination.

## Discussion

The physiological and pathophysiological role of PARP1 in oligodendroglial development and CNS myelinogenesis remains enigmatic. In this study, using genetic and pharmacological approaches applied to *in vivo* and *in vitro* systems, we identify PARP1 as an intrinsic driver for oligodendrocyte differentiation, regeneration, and (re)myelination. PARP1’s PARylation activity plays an essential role in PARP1-regulated oligodendroglial development and myelination. We found that, under physiological conditions, PARP1 PARylates its downstream target proteins that are predominantly involved in RNA metabolism. Our study identified Myef2 as a novel PARylated target, which controls OPC differentiation by PARylation-modulated myelin gene expression. Thus, our studies unravel previously unappreciated functions of PARP1 in oligodendroglial development and CNS (re)myelination and suggest that PARP1-mediated pathway may be a potential therapeutic target for myelin repair in demyelinating disorders.

### PARP1 is required for developmental myelination but not myelin maintenance

PARP1 is highly expressed in the developing CNS and downregulated in adults. Within the oligodendroglial lineage, PARP1 is transiently upregulated in developing OLs and downregulated in fully matured myelinating OLs. These dynamic expression patterns suggest that PARP1 may play a crucial role in developmental myelination. Phenotypic analyses of *Parp1* KO mice indicate that PARP1 is required for oligodendrogenesis and developmental myelination. This conclusion is corroborated by *Olig2:Parp1* cKO mice in which PARP1 is selectively depleted from Olig2-expressing cells commencing from primitive OPCs and by *Pdgfrα:Parp1* cKO mice in which PARP1 is temporally disrupted from neonatal OPCs. These genetic evidences clearly demonstrate an essential role of PARP1 in OPC differentiation and developmental myelination. In contrast to impaired developmental myelination, adult mice of *Parp1* KO or *Parp1* cKO exhibit relatively normal myelin compared with age-matched controls, suggesting that PARP1 plays a minor role in myelin maintenance in the adult CNS. The time-dependent PARP1 function in myelination is in line with its dynamic expression pattern during postnatal CNS development we observed. Interestingly, PARP1 is maintained, albeit at a low level, in all OPCs in the adult CNS, suggesting that PARP1 may regulate the slowly-occurring oligodendrogenesis and myelin turnover in the adult CNS (Yeung et al., 2014; Young et al., 2013) and/or controls the experience-dependent OPC differentiation and adaptive myelination (Mount and Monje, 2017; Xin and Chan, 2020). The defective motor function in the adult *Parp1* KO (fig. S2E-F) and cKO (fig. S3L-M) mice suggests that PARP1 may control other aspects of oligodendroglial functions beyond myelin maintenance. Future studies employing OPC-specific and time-conditional *Parp1* genetic KO models will help address the function of PARP1 in adult OPCs.

### Tightly controlled PARP1 activity plays a minor role in oligodendroglial death

One of the unexpected findings of the current study is the selective and transient activation of PARP1 in OLCs in the CNS white matter during developmental myelination (Fig. 4) and remyelination (Fig. 7A-E). PAR signals are largely confined to the nucleus and barely detected in the cytoplasm of OLCs. The subcellular localization is in line with a major role of PARP1 activity in regulating chromatin structure and gene expression (Gupte et al., 2017). Enzymatically inactive PARP1 acts as a chromatin structural protein, promoting the formation of compact, transcriptionally inactive but poised chromatin (Kim et al., 2004; Tulin and Spradling, 2003). Upon activation, PARP1 and/or its target proteins undergo covalent attachment of the negatively charged PAR chains and subsequent release from compact DNA, facilitating chromatin loosening and gene transcription (Kim et al., 2004; Tulin and Spradling, 2003). PARP1 PARylation activity is tightly controlled by PAR degrading enzyme PARG in OLs, as shown by PARylation accumulation in PARG inhibitor PDD-treated OLs (Fig. 5H), by PARP1 auto-PARylation which provides a negative feedback to PARP1’s enzymatic activity (Gupte et al., 2017; Krishnakumar and Kraus, 2010), and by the downregulation of PARP1 itself in mature OLs (Fig. 1F).

The potential factors that trigger PARP1 activity in developing OLs remain unclear. DNA damage is traditionally thought as the sole activating factor for PARP1 (D’Amours et al., 1999). Consistent with this notion, excessive and sustained PARP1 activation by DNA damaging agents mediates neuronal death in the dish (Alano et al., 2010). However, PARP1 activity in OLs may be unlikely triggered by DNA damage under physiological conditions where the integrity of the genome DNA is maintained. Previous studies suggest that nucleosomes (Kim et al., 2004), ERK1/2 (Kauppinen et al., 2006), and the nuclear NAD^+^ biosynthesizing enzyme NMNAT1 (Ryu et al., 2018; Zhang et al., 2012) could directly stimulate PARP1 enzymatic activity in response to developmental cues. Interestingly, ERK1/2 has been shown to positively regulate developmental myelination (Guardiola-Diaz et al., 2012; Ishii et al., 2013; Ishii et al., 2014; Ishii et al., 2012) and MNNAT1 protects neurons from neurodegenerative insults in the developing CNS (Verghese et al., 2011). In this regard, developmental cue-stimulated PARP1 activity may play a minor role in OL death. In support of this, our data show that disrupting PARP1 and inhibiting its PARylation activity does not affect OL death in the developing CNS. Our results are consistent with previous data showing the PARP1 depletion or inhibition does not affect oligodendroglial cell death induced by peroxynitrite treatment (Scott et al., 2003), suggesting a context-dependent role of PARP1 in cell death/survival (Alano et al., 2010). Furthermore, inhibiting PARP1 activity decreases the number of regenerated OLs under remyelination conditions, suggesting that PARP1 activity is dispensable for OL death during the regenerative process. It remains to be determined whether PARP1 and/or its activity affect OL death or survival under demyelinating condition as previous reported (Veto et al., 2010). Our genetic model of tamoxifen-inducible oligodendroglia-specific PARP1 deletion would be an appropriate tool to answer this question.

### A potential role of PARP1 in regulating RNA metabolism and binding

PARP1 regulates gene expression through its PARylation activity-dependent and independent functions, both of which are not mutually excluded (Gupte et al., 2017). The activity-dependent functions of PARP1 act mainly through PARylating its downstream target proteins. Our results suggest a crucial role of PARP1 in regulating RNA binding and metabolism (Fig. 6), which is consistent with an increasingly appreciated role of PARP1 in RNA biology (Ke et al., 2019). For example, previous studies have shown that PARP1 regulates gene expression at the post-transcriptional level by PARylating and modulating the activity of the polyadenylation polymerase (PAP), an enzyme adding poly(A) tails to protein-coding RNAs (Di Giammartino et al., 2013), the RNA-binding protein HuR (Ke et al., 2017), and heterogeneous nuclear ribonucleoproteins (hnRNPs) (Ji and Tulin, 2013) in a cell- and context-dependent manner. It is worth noting that greater than 86% of the identified PARylated target proteins (108/131) in oligodendroglial lineage cells under the physiological condition are involved in RNA binding (Fig. 6C) and associated with mRNA processing, splicing, translation, and transport (Fig. 6B). Therefore, our proteomic data point to a new direct towards elucidating the molecular mechanism of PARP1 in regulating oligodendroglial differentiation and gene expression.

Interestingly, the extent of MBP reduction at the protein level is greater that at the mRNA level in PARP1 depletion and inhibition assays (Fig. 2O vs Fig. 2P; Fig. 4H vs Fig. 4I). This disproportional reduction suggests that PARP1 may additionally regulate myelin gene expression, directly or indirectly, at the post-transcriptional level. In the current study, we identified a novel PARP1 target protein Myef2 which we showed to control oligodendroglial differentiation at least in part through PARylation-regulated myelin gene expression at the post-transcriptional level (Fig. 6). Myef2 is a transcriptional repressor (van Riel et al., 2012) which had been reported binding to the non-coding strand DNA derived from the *Mbp* promoter and repressing *Mbp* transcription in non-oligodendroglial cell lines (Muralidharan et al., 1997). Myef2 is also an RNA-binding protein harboring three RNA recognition motifs (Keene and Query, 1991; van Riel et al., 2012). We found that PARP1 binds to and PARylates Myef2 in primary oligodendrocytes and that inhibiting PARP1 activity potentiates the binding of Myef2 to myelin gene mRNAs such as *Mbp and Mag,* which repressing myelin protein translation through yet unknown mechanisms. In support of this notion, our functional analysis demonstrates that *Myef2* knockdown promotes myelin gene expression at the post-transcriptional but not mRNA level in primary oligodendrocytes. These data suggest a working model in which PARP1 enzymatic activity promotes oligodendrogenesis by PARylating and releasing the inhibitory effect of Myef2 on myelin protein expression (Fig. 6I). It remains to be determined how Myef2 represses myelin protein expression upon binding to myelin mRNAs and whether Myef2 plays a role in impeding oligodendrocyte differentiation under demyelination and remyelination conditions.

### Targeting PARylation activity to promote oligodendroglial differentiation and regeneration

Our genetic data demonstrate that, among the PARP family, PARP1 is the major PARylating enzyme in the CNS and in the oligodendroglial lineage cells, as evidenced by near absence of PAR in *Parp1* cKO mice (Fig. 3). The mechanisms by which PARP1 regulates oligodendroglial differentiation are likely to be multiple but, at least in part, appear to be through its PARylating activity because inhibiting PARP1’s PARylation activity (without perturbing PARP1 expression) phenocopies the oligodendroglial defects observed in PARP1-deficient primary oligodendroglial cultures and *Parp1* cKO mice. To further elucidate the significance of PARP1-mediated PARylation in oligodendroglial differentiation, we alternatively targeted PARG, the major PAR-degrading enzyme in cells (O’Sullivan et al., 2019), to stabilize PARylation. Inhibiting PARG activity stabilizes cellular PARylation level and accelerates oligodendrogenesis both *in vivo* and *in vitro*, which is further confirmed by *Parg* siRNA-mediated knockdown assays. PARP1/PARG-controlled PARylation are also important in the context of myelin repair, as inhibiting PARylation negatively impact oligodendrocyte regeneration and remyelination in a toxin-induced demyelination model (Fig. 7). All our data suggest that PARP1/PARG-controlled PARylation activity may be a potential therapeutic target for myelin regeneration.

Interestingly, *in vivo* PARG inhibition by PDD treatment increases PAR signal in Sox10^+^ OLCs and does not lead to massive ectopic PAR accumulation in non-oligodendroglial lineage cells (Fig. 5C). This observation suggests that, despite ubiquitous expression of PARG, the de-PARylation activity of PARG exists in cells simultaneously exhibiting PARP1’s PARylation activities, which are identified as OLCs (Fig. 4). This finding also implies that the cellular PARylation level is tightly controlled by the cooperative activity of PARP1 and PARG in OLCs. A previous study showed that germinal PARG depletion results in very early embryonic lethality due to aberrant PARylation accumulation and massive apoptosis of the blastocytes (Koh et al., 2004). Our *in vitro* data showed that a 40-60% knockdown of PARG is sufficient to accelerate oligodendroglial differentiation and upregulate the potent oligodendroglial pro-differentiation factor Myrf (Emery et al., 2009). It would be very interesting to study whether reducing or inhibiting PARG could potentially promote OL regeneration in demyelinating animal models.

### PARP1-mediated PARylation is relevant in demyelinating and neurodegenerative disorders

The induction of PARP1 and PARylation has been observed in OLCs in the active lesions of multiple sclerosis (MS) with much less being seen in the chronic lesions (Veto et al., 2010), suggesting that intervening PARylation activity may be clinically relevant for oligodendrocyte regeneration and myelin repair in MS patients. However, it is unclear whether oligodendroglial PARP1 in MS lesions is a sign of remyelination failure or an indication of remyelination occurrence. PARP1 and its activity are induced in oligodendroglial lineage cells in the active lesions of MS brains where extensive spontaneous remyelination occurs, suggesting PARP1 as a potential target for myelin repair. Our data suggest the induction of PARP1 and/or PARylation in MS active lesions may be an indication of spontaneous remyelination occurrence. Consistently, our genetic and pharmacological studies of developmental myelination and remyelination suggest that augmenting oligodendroglial PARylation may be beneficial in myelin repair. Recently, neuronal PARylation has been reported to potentiate neurodegeneration in animal models of Parkinson’s disease (Kam et al., 2018), again suggesting that PARP1-mediated PARylation may play a cell type-dependent role in disease pathology. Thus, cell type-specific strategies are needed to elucidate the detrimental or beneficial role of cellular PARylation in demyelinating disorders and other neurodegenerative diseases. Our *Cre-loxP* animal models would help address these important issues in future studies.

## Materials and Methods

### Animals

The following transgenic mice were used in our study: *Parp1* knockout mice (RRID: IMSR_JAX:002779) and *Parp1-floxed* mice provided by Dr. Kraus (Luo et al., 2017). *Olig2-Cre* (RRID: IMSR_JAX:025567), *Pdgfrα-CreER^T2^* (RRID: IMSR_ JAX:018280) and *Rosa26-LoxP-STOP-LoxP-*EYFP (referred to as *Rosa26-EYFP*) (RRID: IMSR_JAX:006148) were purchased from the Jackson Laboratory. *Cre* transgene was always maintained as heterozygosity. We crossed *Cre* lines with *Parp1^fl/fl^* mice to generate *Parp1* cKO mice. For PARylation study, neonatal mice were injected with PARP inhibitor 4-hydroxyquinazoline (4HQ, H57807, Sigma-Aldrich) 100 mg/kg body weight, PARG inhibitor PDD 00017273 (HY-108360, MedChemExpress) 10 gm/kg body weight or vehicle control *subcutaneously* from postnatal day 2 (P2) to P10, and euthanized for analysis 2 h after the last injection. For postnatal studies, the day when pups were born was designated as P0. Both male and female mice were used in this study. All animals were from C57BL/6 background and maintained in 12 h light/dark cycle with water and food. Animals and procedures in this study were approved by the Institutional Animal Care and Use Committee at the University of California, Davis.

### Gene conditional KO by tamoxifen treatment

Tamoxifen (T5648, Sigma-Aldrich) was dissolved in a mixture of ethanol and sunflower seed oil (1:9, v/v) at the concentration of 30 mg/ml. Tamoxifen was administrated subcutaneously to neonatal pups (*Pdgfrα-CreER^T2^:Parp1^fl/fl^* and littermate controls) on P1, P2, and P3 daily at a dose of 10 μl (300 μg) once a day and mice were euthanized at P9 according to our previous protocols (Zhang et al., 2020b; Zhang et al., 2018b).

### Tissue preparation and immunohistochemistry (IHC)

Tissue preparation and IHC were conducted as previously described (Zhang et al., 2020a). After anesthetization by ketamine/xylazine mixture, mice were transcardially perfused with ice-cold phosphate buffered saline (PBS, pH = 7.0, BP399-20, Fisher Chemical). Tissues were collected, and immediately placed on dry ice for protein or RNA extraction or fixed in fresh 4% paraformaldehyde (PFA, #1570-S, Electron Microscopy Science) for histological study. After post-fix in 4% PFA for 2 h at room temperature (RT), tissues were washed with PBS three times, 15 min each time, cryopreserved in 30% sucrose (S5-3, Fisher Chemical) in PBS overnight at 4°C and embedded in OCT (VWR International, Wetzlar, Germany). Serial coronal sections (12 μm) were cut by a Leica Cryostat (CM 1900-3-1) and stored in −80 °C. IHC was conducted as below: slices were air dry at RT for 2 h, and blocked with 10% donkey serum in 0.1% Triton X-100/PBS (v/v) for 1 h at RT, followed by incubation with primary antibodies overnight at 4°C. After wash with PBST (PBS with 0.1% Tween-20, v/v), slices were incubated with fluorescence conjugated secondary antibodies for 2 h at RT. DAPI was applied as nuclear counterstain. All images shown were obtained by Nikon A1 confocal microscope. 10 μm of optical thickness sections were obtained by confocal z-stacking (step size 1 μm) and projected into one flattened image for quantification. The following antibodies were used in IHC: PARP1 (1:100, 39559, RRID: AB_2793257, Active Motif; 1:100, 66520-1-Ig, RRID: AB_2881883, Proteintech), Anti-poly-ADP-ribose binding reagent (1:100, MABE1031, RRID: AB_2665467, Millipore), Sox2 (1:500, sc-17320, RRID: AB_2286684 Santa Cruz Biotechnology), CC1 (1:200, OP80, RRID: AB_213434, Calbiochem), TCF7l2 (1:100, 2569S, RRID: AB_2199816, Cell Signaling Technology; 05-511, RRID: AB_309772, Millipore), PDGFRα (1:100, AF1062, RRID: AB_2236897, R&D System), EYFP (1:200, 06-896, RRID: AB_310288, Millipore), Sox10 (1:100, ab27655, RRID: AB_778021, Abcam), MBP (1:100, NB600-717, RRID: AB_2139899, Novus), Myef2 (1:100, 16051-1-AP, RRID: AB_2146869, Proteintech), SMI312 (1:100, RRID: AB_2565383, 837801, BioLegend), Iba1 (1:100, NB100-1028, RRID: AB_521594, Novus), GFAP (1:100, MAB360, RRID: AB_2275425, Millipore). All secondary antibodies DyLight 488- or DyLight549-conjugated (Fab)2 fragments (1:500) were from Jackson ImmunoResearch Laboratories.

### Black-Gold II myelin staining

Black-Gold II myelin stain (AG105, Millipore) was performed according to the manufacturer’s instruction. The fixed, frozen sections (12 um) were incubated with 0.3% Black-Gold II stain for 20 min at 60°C. The stain was fixed with 1% sodium thiosulfate solution, and then washed, dehydrated using a series of gradated alcohol (50%, 75%, 85%, 95% and 100%), cleared in xylene, and coverslipped with mounting medium (SP15-100, Fisher Scientific). Staining was visualized using an Olympus BX61 microscope and quantified by ImageJ software.

### Toluidine blue staining and transmission electron microscopy (TEM)

Tissue processing for semithin sections of toluidine blue staining and ultrathin sections of TEM was performed as previously described with some modifications [4,6]. In brief, mice were anesthetized with ketamine/xylazine mixture and perfused with 4% PFA, followed by 3% glutaraldehyde (16130, Electron Microscopy Science, dilute in PBS, pH 7.4) at a speed of 5 ml per minute. Spinal cord was carefully dissected out and fixed with 3% glutaraldehyde overnight, followed by wash with 0.2 M sodium cacodylate buffer (pH 7.2, #11653, Electron Microscopy Science) twice, 10 min each time, post-fixed with 2% (w/v) aqueous osmium tetroxide (#19152, Electron Microscopy Science) for 2 h and washed with sodium cacodylate twice, 10 min each time. The resulting spinal cord was then dehydrated with 50, 70, 90, and 100% ethanol, followed by wash in propylene oxide three times, 30 min each time, and incubation with 1:1 mixture of Propylene Oxide:Eponate Resin overnight, and with 1:3 mixture of Propylene Oxide:Eponate Resin for 10 h, and with 100% Eponate Resin overnight. The resulting specimens were embedded in EMBed-812 Resin for 2 days at 65°C. Semithin (500 nm) sections were cut by using a Leica EM UC6 microtome and then incubated with 2% toluidine blue (#19451, Ted Pella Inc.) at 100°C for 2 min, followed by imaging on Olympus BX61 microscope. Ultrathin (70 - 80 nm) sections were cut on a Leica EM UC7 microtome and collected on 1 mm Formvar-coated copper slot grids, double stained with uranyl acetate and lead citrate, followed by imaging on a CM120 electron microscope.

### Primary rodent oligodendrocyte progenitor cells (OPCs) culture, differentiation, and treatment

Primary OPCs were isolated from cerebral cortices of P0 to P2 mice using immune-panning procedure according to our previous protocol [7,8]. After stripping the meninges, cerebral cortices were enzymatically digested using papain (20 U/ml, LK003176, Worthington) supplemented with DNase I (250 U/ml; D5025, Sigma) and D-(+)-glucose (0.36%; 0188, AMRESCO), and mechanically triturated to obtain single cell. The cell suspension was centrifuged, resuspended in DMEM medium (1196092, Thermo Fisher) with 10% heat-inactivated fetal bovine serum (12306-C, Sigma) and penicillin/streptomycin (P/S, 15140122, Thermo Fisher), and plated on poly-D-lysine (PDL, A003-E, Millipore) coated 10 cm dishes. After 24 h incubation, cells were washed using HBSS (with calcium and magnesium, 24020117, Thermo Fisher), and cultured in serum-free growth medium (GM), containing 30% of B104 neuroblastoma medium and 70% of N1 medium (DMEM with 5 μg/ml insulin (I6634, Sigma), 50 μg/ml apo-transferrin (T2036, Sigma), 100 μM putrescine (P5780, Sigma), 30 nM Sodium selenite (S5261, Sigma), 20 nM progesterone (#P0130, Sigma) until 80% of confluency. The mixed glial cells were then dissociated in single cell suspension, incubated on Thy1.2 (CD90.2) antibody (105302, Biolegend) coated Petri-dish to deplete astrocytes, neurons and meningeal cells and then seeded on NG2 antibody (#AB5320, Millipore) coated Petri-dish to select OPCs. The isolated OPCs were then cultured on PDL-coated plates using GM plus 5 ng/ml FGF (450-33, Peprotech), 4 ng/ml PDGF-AA (315-17, Peprotech), 50 µM Forskolin (6652995, Peprotech,) and glutamax (35050, Thermo Fisher). To induce differentiation, OPCs were cultured in the differentiation medium (DM), consisting of F12/high-glucose DMEM (11330032, Thermo Fisher Scientific) plus 12.5 μg/ml insulin, 100 μM Putrescine, 24 nM Sodium selenite, 10 nM Progesterone, 10 ng/ml Biotin, 50 μg/ml Transferrin (T8158, Sigma), 30 ng/ml 3,3′,5-Triiodo-L-thyronine (T5516, Sigma), 40 ng/ml L-Thyroxine (T0397, Sigma-Aldrich), glutamax and P/S. OPCs with 80% confluence were treated with 10 μM 4HQ, 1 μM PDD or vehicle control in the DM medium. OPCs differentiation was analyzed at the time points indicated in Figure 4G and 5G.

### Immunocytochemistry (ICC)

Cells cultured on glass slides were fixed in 4 % PFA for 30 min, permeabilized with 0.1 % Triton X-100 in PBS and blocked with 10 % donkey serum. The cells were then washed with PBS and incubated with Primary antibodies overnight at 4 °C, followed by fluorescence-conjugated secondary antibodies (1:200) for 2 h at room temperature. Nuclei were visualized using DAPI. Fluorescent images were observed with Nikon A1 confocal microscope. Intensity of MBP fluorescence were quantified using ImageJ software.

### Cell viability assay

The cell viability was measured using 3-(4, 5-dimethylthiazol-2-yl)-2, 5-diphenyl tetrazolium bromide (MTT, M5655, Sigma) as described before [10]. Cells were seeded into 48-well plates at a density of 1 × 10^5^ cells/mL and cultured for 24 h before further treatment. 200 μl of MTT solution (5 mg/mL, diluted in PBS) was added to each well and incubated at 37 °C for 4 h. The medium was removed and 200 μl of DMSO was added to each well to dissolve formazan crystals. Optical densities (OD) were determined by a microplate reader (SpectraMax i3x, Molecular Devices) at 570 nm and 650 nm. Cell viability was expressed as a percentage with the control cells, which was taken as 100%.

### Protein extraction and Western blot assay

Protein extraction and Western blot assay were performed as previously described with some modifications (Tabrizi et al.). Tissue or cells were lysed in N-PER™ Neuronal Protein Extraction Reagent (#87792, Thermo Fisher) supplemented with protease and phosphatase inhibitor cocktail (#78440, Thermo Fisher) and PMSF (#8553S, Cell Signaling Technology). After incubation on ice for 10 min and centrifugation at 10,000 × g for 10 min at 4°C, the concentrations of each samples were measured using BCA protein assay kit (#23225, Thermo Fisher Scientific). Equal volumes (30 μg) of cell lysates from each condition were resolved by AnykD Mini-PROTEAN TGX precast gels (#456-9035, BIO-RAD) or 7.5% Mini-PROTEAN TGX precast gels (#456-8024, BIO-RAD). The proteins were transferred onto 0.2 μm nitrocellulose membrane (#1704158, BIO-RAD) using Trans-blot Turbo Transfer system (#1704150, BIO-RAD). After blocking with 5% BSA (#9998, Cell signaling) for 1 h at room temperature and the membranes were incubated with primary antibodies overnight at 4 °C, followed by suitable HRP-conjugated secondary antibodies. Proteins of interest were visualized by Western Lightening Plus ECL (NEL103001EA, Perkin Elmer). NIH Image J was used to analyze protein levels. Primary and secondary antibodies used were: PARP1 (1:1000, 39559, RRID: AB_2793257, Active Motif), Anti-poly-ADP-ribose binding reagent (1:1000, MABE1031, RRID: AB_2665467, Millipore), Sox2 (1:1000, sc-17320, RRID: AB_2286684 Santa Cruz Biotechnology), TCF7l2 (1:100, 2569S, RRID: AB_2199816, Cell Signaling Technology; 05-511, RRID: AB_309772, Millipore), Myef2 (1:100, 16051-1-AP, RRID: AB_2146869, Proteintech), MBP (1:100, NB600-717, RRID: AB_2139899, Novus), MAG (1:1000, AB1567, RRID: AB_2110397, Millipore), PLP (1:1000, PA3-151, RRID: AB_2165785, Thermo Fisher Scientific), TLE3 (1:1000, #11372-1-AP, RRID: AB_2203743, Proteintech), AIF (1:1000, ab1998, RRID: AB_302748, Abcam), Gasdermin D (1:1000, 93709, RRID: AB_2800210, Cell Signaling Technology), MLKL (1:1000, Cat# 66675-1-Ig, RRID: AB_2882029, Proteintech), p-MLKL (S345) (1:1000, D6E3G, RRID: AB_2799112, Cell Signaling Technology), β-actin (1:1000, #4967L, RRID: AB_330288, Cell Signaling Technology), and HRP goat anti-rabbit (1:3000, 31460, RRID: AB_228341, Thermo Fisher Scientific), anti-mouse (1:3000, 31430, RRID: AB_228307, Thermo Fisher Scientific) or anti-rat (1:3000, #7077, RRID: AB_954646, Cell Signaling Technology) secondary antibodies.

### Co-Immunoprecipitation (Co-IP)

For Western blotting assay, Co-IP was performed with Pierce Crosslink Magnetic IP/Co-IP kit (#88805, Thermo Fisher) following the manufacturer’s instructions. 10 μg of primary antibodies or isotype control IgG were covalently cross-linked to 25 μl of protein A/G magnetic beads (#88802, Thermo Fisher). Proteins were extracted with the Pierce IP Lysis/Wash Buffer (#87787, Thermo Fisher) supplemented with PMSF (#8553S, Cell Signaling Technology) and protease and phosphatase inhibitor cocktail (#78440, Thermo Fisher). A portion of each sample was saved as input. An equal amount (1 mg) of each protein extract was incubated with protein A/G magnetic beads cross-linked with primary antibodies or isotype control IgG overnight at 4°C. The beads were then washed to remove non-bound material and protein was eluted in a low-PH elution buffer that dissociates bound antigen from the antibody-crosslinked beads. The elute was then added with Neutralization buffer to neutralize the low PH and Lane marker sample buffer containing β-mercaptoethanol for SDS-PAGE and Western blotting. Primary antibodies and isotype control IgG used were: PARP1 (39559, RRID: AB_2793257, Active Motif), PAR (4335-MC-100, RRID: AB_2572318, Trevigen), Sox2 (sc-17320, RRID: AB_2286684 Santa Cruz Biotechnology), Myef2 (16051-1-AP, RRID: AB_2146869, Proteintech), Normal rabbit IgG (#2729, RRID: AB_1031062, Cell Signaling Technology), Normal goat IgG (AB-108-C, RRID: AB_354267, R&D Systems).

For LC-MS/MS assay, Co-IP was performed with Pierce Classic Magnetic IP/Co-IP kit (#88804, Thermo Fisher) according to the instructions of manufacturer. 1 mg of protein lysate was incubated with 10 μg of primary antibodies or isotype control IgG overnight at 4°C. Pierce Protein A/G Magnetic Beads were washed three times using Pierce IP Lysis/Wash Buffer for LC-MS/MS analysis.

### LC-MS/MS

For protein digestion, samples on magnetic beads were washed four times with 200 ul of 50mM ammonium bicarbonate (AMBIC) with a 20 min shake time at 4°C. 2.5 ug of trypsin gold (Mass spectrometry grade, V528A, Promega) was added to the beads and samples were digested overnight at 800 rpm shake speed at RT. After overnight digestion, the peptide extracts were reduced in volume by vacuum centrifugation and a small portion of the extract was used for fluorometric peptide quantification (Thermo scientific Pierce). Samples were analyzed for LC-MS/MS analysis by UC Davis Proteomics core. One microgram of sample based on the fluorometric peptide assay was loaded for each LC-MS/MS on a Thermo Scientific Q Exactive Plus Orbitrap Mass spectrometer in conjunction Proxeon Easy-nLC II HPLC (Thermo Scientific) and Proxeon nanospray source. Tandem mass spectra were extracted and charge state deconvoluted by Proteome Discoverer (Thermo Scientific) All MS/MS samples were analyzed using X! Tandem (The GPM, thegpm.org; version X! Tandem Alanine (2017.2.1.4)). Scaffold (version Scaffold_4.8.4, Proteome Software Inc., Portland, OR) was used to validate MS/MS based peptide and protein identifications.

### RNA extraction and quantitative real-time PCR (qRT-PCR)

RNA isolation was performed using the RNeasy Lipid Tissue Mini Kit (#74804, QIAGEN) with RNase-Free DNase Set (#79254, QIAGEN) to remove genomic DNA [11]. RNA concentration was measured by Nanodrop 2000 Spectrophotometer (Thermo Fisher Scientific). cDNA was synthesized by Qiagen Omniscript RT Kit (#205111, Qiagen). RT-qPCR was performed using QuantiTect SYBR® Green PCR Kit (#204145, QIAGEN) on Agilent MP3005P thermocycler. For quantification, the mRNA expression level of interested genes in each sample was normalized to that of the internal control gene Hsp90, and fold change in gene expression was calculated based on the equation 2^(Ct(cycle threshold) of Hsp90 − Ct of indicated genes). The gene expression levels in control groups were normalized to 1. The qPCR primers used in this study were *Parp1* (F: AAGGCGGAGAAGACATTGGG, R: ACCATCTTCTTGGACAGGCG), *Parp2* (F: TGGAAGGCGAGTGCTAAATGA, R: GGGCTTTGCCCTTTAACAGC), *Parp3* (F: AAGGCCTTATCTCCCCAGGT, R: TTCACATCCAGGTTCATGAGGG), *Sox10* (F: ACACCTTGGGACACGGTTTTC, R: TAGGTCTTGTTCCTCGGCCAT), Pdgfr*α* (F: GAGAACAACGGAGGAGC, R: GCTGAGGACCAGAAAGACC) *Enpp6* (F: CAGAGAGATTGTGA ACAGAGGC, R: CCGATCATCTGGTGGACCT), *Mbp* (F: GGCGGTGACAGACT CCAAG, R: GAAGCTCGTCGGACTCTGAG), *Mag* (F: CTGCCGCTGTTTTGGATAATGA, R: CATCGGGG AAGTCGAAACGG), *Mog* (F: AGCTGCTT CCTCTCCCTTCTC, R: ACTAAAGCCCGGATGGGATAC), *Plp-E3b* (targeting the exon 3b of Plp1 gene, which is only expressed in myelinating OLs) (F: GTTCCAGAGGCCAACATCAAG, R: CTTGTCGGGATGTCCTAGCC), *Myrf* (F: CAGACCCAGGTGCTACAC, R: TCCTGCTTGATCATTCCGTTC), *Bmp4*: (F: TTCCTGGTAACCGAATGCTGA, R: CCTGAATCTCGGCGACTTTTT), *Tcf7l2* (F: GGAGGAGAAGAACTCGGAAAA, R: ATCGGAGG AGCTGTTTTGATT), *Gfap* (GTGTCAGAAGGCCACCTCAAG, R: CGAGTCCTTAATGACCTCACCAT), *Bcas1* (F: AGAAGCGAAAGGCTCGGAAG, R: AGGGACAG AATAACTCAGAGTGT), *Cnp* (F: TTTACCCGCAAAAGCCACACA, R: CACCGTGT CCTCATCTTGAAG), *Qk* or *CC1* (F: CTGGACGA AGAAATTAGCAGAGT, R: ACTGCCATTTAACG TGTCATTGT), *Ugt8a* (F: GAACATGGCTTTGTCCTGGT, R: CATGGCTTAGGAAGGCTCTG), *Mobp* (F: AACTCCAAGCGTGAGATCGT, R: - CAGAGGCTGTCCATTCACAA), *Grp17* (F: ACACAGTTGTCTGCCTGCAA, R: TGACCGTGGTGATGAATGGG), *Aqp4* (F: ATCAGCATCGCTAAGTCCGTC, R: GAGGTGTGACCAGGTAGAGGA), *Aldh1l1* (F: AGCCACCTATGAGGGCATTC, R: TGAGTGTCGAGTTGAAAAACGTC), *Parg* (F: AACGCCACCTCGTTTGTTTTC, R: CACAGAACTCATCATGGAGTCAA), *Cd68* (F: TGTCTGATCTTGCTAGGACCG, R: GAGAGTAACGGCCTTTTTGTGA), *Vim* (F: GAGGAGATGAGGGAGTTGCG, R: CTGCAATTTTTCTCGCAGCC), *Aif1* (F: GGATCAACAAGCAATTCCTCGA, R: CTGAGAAAGTCAGAGTAGCTGA), *Myef2*- (F: CGGGTTTTGGAGGAGTGAATAG, R: CAATATCACCACGTCCGAAATCT), *Hsp90* (F: AAACAAGGAGATTT TCCTCCGC, R: CCGTCAGGCTCTCATATCGAAT), *Parg*-rat (F: TAAGGACGCTCCCGTACAGT, R: TTGAAAACAAACGAGGTGACG), *Myef2*-rat (F: GGAATGGATGGTCCGGGTTT, R: CGCGGTACAACTCTCCCATT), *Mbp*-rat (F: TTGACTCCATCGGGCGCTTCTTTA, R: TTCATCTTGGGTCCTCTGCGACTT) *β-actin*-rat: (F: CGTCTTCCCCTCCATCGT, R: GGAGTCCTTCTGACCCATACC)

### RNA-sequencing (RNA-seq) and bioinformatic analysis

Total RNA was prepared from forebrains of *Pdgfrα-CreER^T2^*, *Parp1^fl/fl^* mice (n = 3) and controls (n = 3) using QIAGEN RNeasy for lipid tissues (#74804) with on-column DNase I digestion. The quality of RNA samples was determined by the Bioanalyzer 2100 system (Agilent Technologies). The RNA integrity number (RIN) of all our RNA samples used for RNA-seq and qPCR was greater than 6.8. The cDNA library was prepared using the NEBNext Ultra Directional RNA Library Prep Kit (#E7420, New England BioLabs) for Illumina, and sequenced on the Illumina HiSeq 4000 sequencing platform. Single-end clean reads were aligned to the reference genome (mouse genome mm10) using TopHat version 2.0.12. Differentially expressed genes were analyzed using DESeq version 1.10.1, and *p* < 0.05 was assigned as differentially expressed genes (DEGs). Gene ontology (GO) enrichment analysis of DEGs was performed using the National Institutes of Health online tool DAVID (https://david.ncifcrf.gov/). GO terms with adjusted P value less than 0.05 were considered significantly downregulated.

### Primary rodent OPCs culture

Primary mixed glial cells were prepared from P0-P2 rat cerebral cortices as previous described (Zhang et al., 2020a). Mixed glial cells were plated on PDL-coated T75 flasks in high glucose DMEM medium (Gibco,11965-092) supplemented with 10% heat-inactivated fetal bovine serum and 1% Penicillin/Streptomycin. Medium was changed every 2 days until astrocytes are confluent. Flasks were shaken for 1 hour at 37°C and 200 rpm on an orbital shaker (Cat# C491, Hanchen) to remove microglia. After wash with PBS, 20 ml 10% FBS/DMEM was added into each flask, followed by shaking for 6 h at 37°C and 200 rpm to get OPCs. OPCs were seeded on PDL-coated plates in GM for further study.

### RNA immunoprecipitation (RIP)

RIP was conducted as reported. D2 OLs were lysed in Pierce IP Lysis/Wash Buffer supplemented with protease inhibitor (5871, Cell Signaling Technology) and 100 U/ml RNAseOUT (10777019, Thermo Fisher) for 30 min on ice. After centrifugation, the protein lysate was incubated with 10 μg of Myef2, or normal rabbit IgG antibody overnight at 4°C. 50 μl of protein G Dynabeads (10003D, Life Technologies) were added to the lysate and antibody mix and incubated for 2 h at 4°C. The beads were then washed five times using 1 ml of lysis buffer. RNA binding to Myef2 protein and 10% of the input lysate were extracted by Buffer RLT supplemented with β-mercaptoethanol (10 μl per 1 ml Buffer RLT) and vortexed to mix. 1 volume of 70% ethanol was added to the homogenized lysate, and sample was transferred to RNeasy Mini spin column, centrifuged and washed with Buffer RW1 and Buffer RPE. RNA was eluted by 25 μl of RNase-free water. Purified RNA was then analyzed by RT-qPCR. *<u>Mbp</u>* primers (F: TGGAAGGATGGTCAGTAGGG, R: TTTAACCTCCTCTCCCAACTCT). *Mag* primers (F: ACAGCGTCCTGGACATCATCAACA, R: ATGCAGCTGACCTCTACTTCCGTT).

### siRNA transfection

The siRNA oligonucleotides used to knockdown *Parg* (sense: 5’-GACAUUAACUUCAAUCGGU-3’, antisense: 5’-ACCGAUUGAAGUUAAUGUC-3’), Myef2 (sense: 5’-CUUUGGUGGAGUUGGCCGA-3’, antisense: 5’-UCGGCCAACUCCACCAAAG-3’) or siRNA universal negative control (SIC001, Sigma-Aldrich) were purchased from Sigma-Aldrich. Primary OPCs growing on 6-well plates at 80% confluency were transfected with a concentration of 25 nM siRNA and 9 μl HiPerFect transfection reagent (301705, Qiagen) in the differentiation medium. Cells were incubated with the medium containing siRNA at the time points indicated in Figure 5M and 6J. Cells were processed for analysis after transfection.

### Cuprizone model

Experimental demyelination was induced by feeding 8-week-old male mice with 0.2% cuprizone (TD.140800, Envigo) diet. After 6 weeks of treatment, cuprizone feedings were discontinued and changed to a regular diet for 1 week (6+1 W) or 6 weeks (6+6 W) for remyelination. To study the effect on remyelination, 4HQ (100 mg/kg) or DMSO vehicle were intraperitoneally injected daily starting from day 1 through day 7 after returning to normal diet, and mice in each group were euthanized at weeks 7 (6+1 W) for further analysis.

### Rotarod test

Accelerating rotarod test was used for motor function assessment. The starting speed of rotarod was 4 rotations per minute (rpm) and the maximal speed was 40 rpm with 1.2 rpm incremental every 10 s. Mice were trained for two consecutive days (4 trials each day with 60 min interval between trials) followed by data collection on the third day. The retention time of each mouse on rod was recorded and calculated by averaging 3 individual trials.

### Statistical analyses

Quantification was performed by observers blind to genotype and treatment. Data were presented as mean ± s.e.m. except stated otherwise. Scatter dot plots were used to quantify data throughout our manuscript. Each dot in the scatter dot plots represents one mouse or one independent experiment. Shapiro-Wilk approach was used for testing data normality. Unpaired two-tailed Student’s t test was used for statistically analyzing two groups of data and degree of freedom (df) were presented as t_(df)_ in figure legends. One-way ANOVA followed by Tukey’s post-test was used for statistically analyzing three or more groups of data. The F ratio, and DFn and DFd was presented as F(DFn, DFd) in the figure legends where DFn stands for degree of freedom numerator and DFd for degree of freedom of denominator. All data graphing and statistical analyses were performed using GraphPad Prism version 8.0. P value less than 0.05 was considered as significant. ns stands for not significant with P value greater than 0.05.

## Acknowledgments

We thank W. Lee Kraus and Xin Luo at UT Southwestern for generously providing *Parp1*-floxed mice. This work is funded in part by NIH/NINDS (R21NS109790 and R01NS094559) and by Shriners Hospitals for Children (85107-NCA-19, 84553-NCA-18, and 84307-NCAL).

## Supplementary Materials

**fig. S1.**
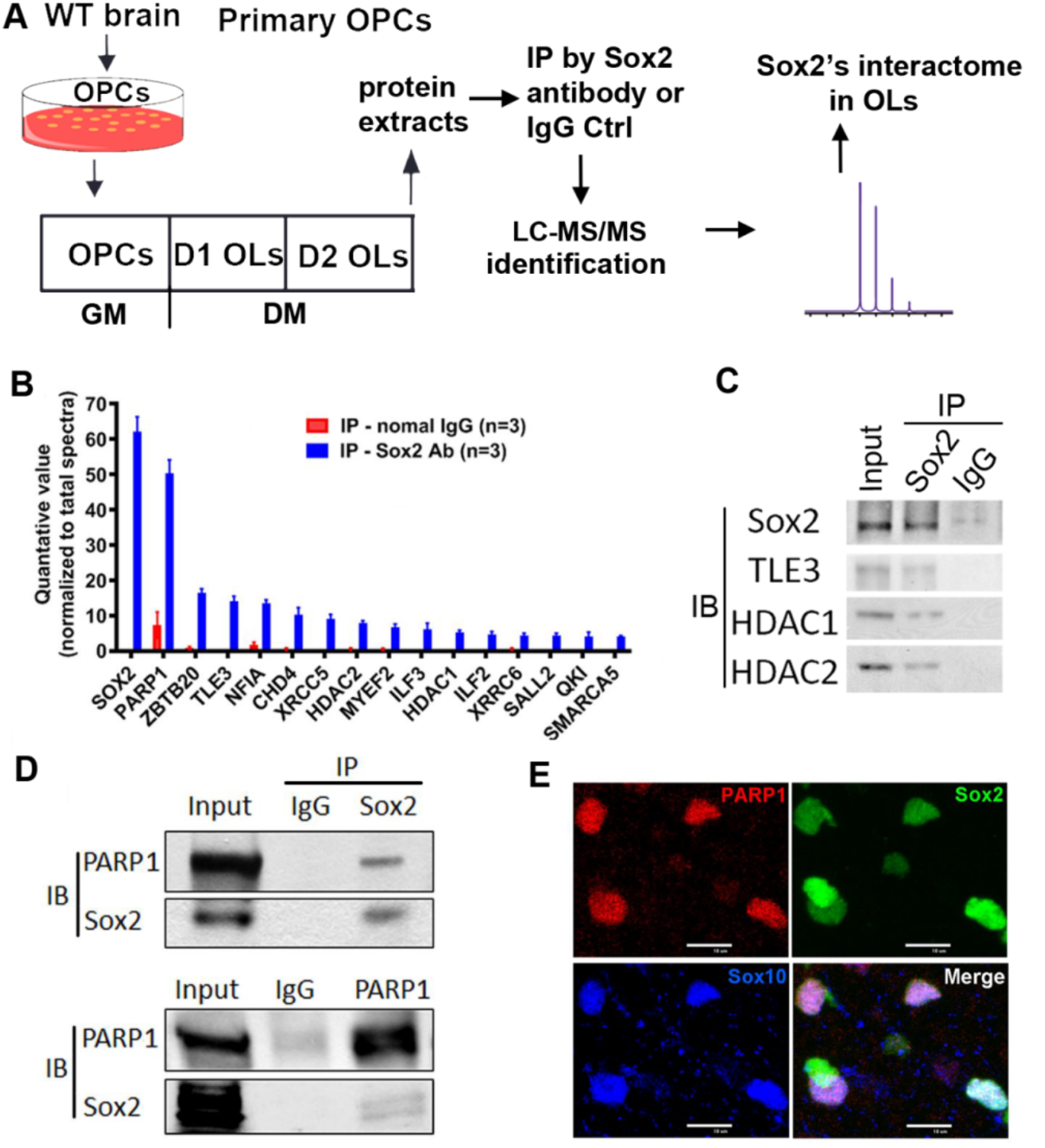
Identification of PARP1 as a SOX2 binding partner. **A**, flow chart of identifying Sox2’s binding partner at the proteomic level in oligodendrocytes (OLs). Protein extracts from Day 2 differentiated primary oligodendrocytes (D2 OLs) were immunoprecipitated (IP) by anti-Sox2 antibody and were identified by liquid chromatography coupled with tandem mass spectrometry (LC-MS/MS). Anti-rabbit IgG antibody was used as negative control. n = 3 each group. GM, growth medium, DM, differentiation medium. Our previous studies have reported that primary OLs at ∼ Day 2 differentiation are newly formed OLs and those at ∼D4 are sheath-forming mature OLs. **B**, representative candidates of Sox2 binding proteins identified from D2 primary OLs. **C**, confirmation of representative candidates of Sox2’s binding partners (TLE3, HDAC1/2) identified from LC-MS/MS by Western blot (IB) in D2 primary OLs. **D,** confirmation of Sox2 and PARP1 interaction in D2 primary OLs. **E,** triple immunohistochemistry (IHC) staining of PARP1, Sox2 and pan-oligodendroglial lineage marker Sox10 in the white matter of mouse spinal cord at postnatal day 7. Scale bar = 10μm.

**fig. S2.**
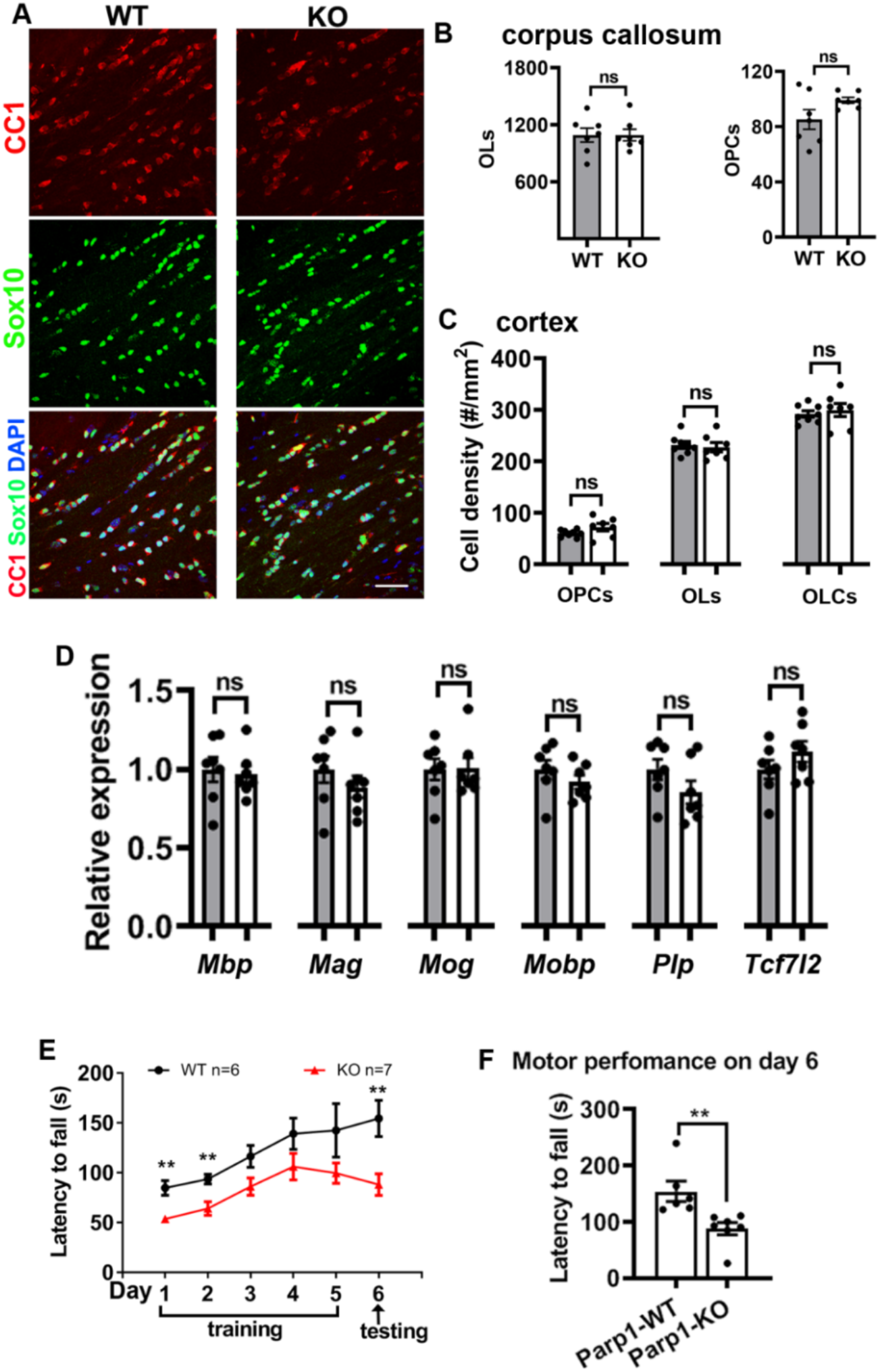
Normal oligodendrocyte generation and myelin gene expression and impaired motor performance of adult *Parp1* KO mice. **A**, representative confocal images of Sox10 and CC1 in the corpus callosum of P70 mice. Scale bar = 50μm. **B-C**, densities (# per mm^2^) of Sox10^+^CC1^+^ OLs, Sox10^+^CC1^-^ OPCs, and Sox10^+^ oligodendroglial lineage cells (OLCs) in the corpus callosum (**B,** n = 7 WT and 7 KO) and the cingulate cortex (**C,** n = 8 WT and 7 KO). **D**, qRT-PCR assay for myelin-specific genes and pre-myelinating OL marker gene *Tcf7l2* in the brain. N = 7 WT, 7 KO. **E**. motor training and testing by accelerating Rotarod. Adult (P60) *Parp1* WT and KO mice were training for retaining on the accelerating rod for 5 consecutive days (3 trials per day) and motor performance was probed at day 6. The time on the rod prior to fall (latency to fall, seconds) was recorded. Rotarod testing was performed according to our previous protocols. **F**, motor performance on day 6. Statistic methods: unpaired two-tailed Student’s *t* test (B-F) and two-way ANOVA followed by Sidak multiple comparison test (E). \**P* < 0.05; \*\**P* < 0.01; \*\*\**P* < 0.001, ns, not significant. **B**, OLs *t*_(12)_ = 0.0007, *P* = 0.9995; OPCs Welch-corrected *t*_(7.067)_ = 1.890, *P* = 0.1002. **C**, OPCs *t*_(13)_ = 0.3897, *P* = 0.7030; OLs *t*_(13)_ = 0.5576, *P* = 0.5866; OLCs Welch-corrected *t*_(7.406)_ = 1.548, *P* = 0.1633. **D**, Mbp *t*_(12)_ = 0.3286, *P* = 0.7481; Mag *t*_(12)_ = 1.013, *P* = 0.3309; Mog *t*_(12)_ = 0.05157, *P* = 0.9598; Mobp *t*_(12)_ = 1.073, *P* = 0.3044; Plp *t*_(12)_ = 1.489, *P* = 0.1623; Tcf7l2 *t*_(12)_ = 1.271, *P* = 0.2278. **E**, genotype *F*_(1, 66)_ = 28.47 *P* < 0.0001; time-course *F*_(5, 66)_ = 6.959 *P* < 0.0001. **F**, *t*_(11)_ = 3.247, *P* = 0.0078.

**fig. S3.**
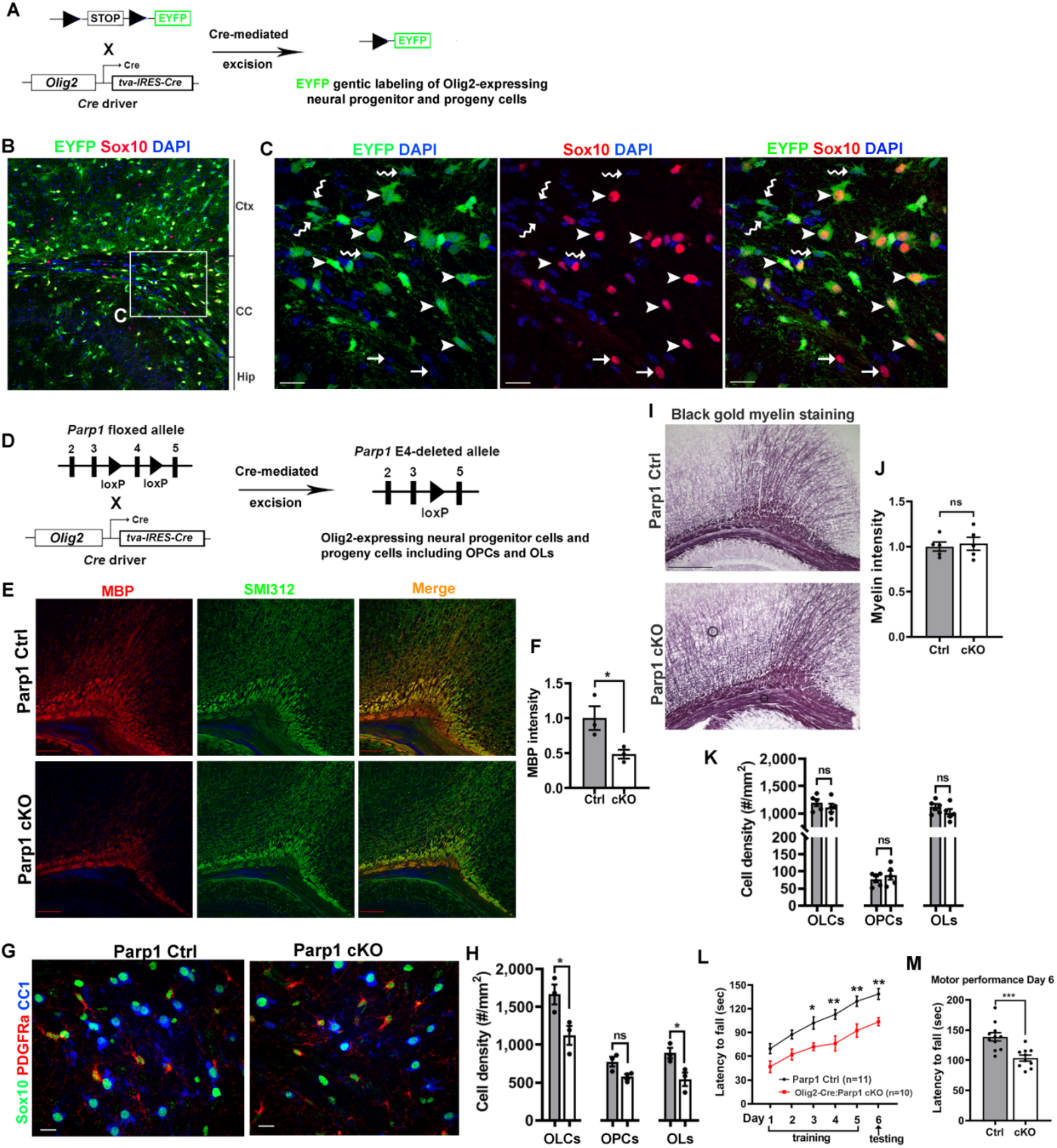
PARP1 depletion in Olig2-expressing neural progenitor cells inhibits OL differentiation during postnatal CNS development. **A**, fate-mapping strategy of Olig2-expressing cells. Constitutive *Olig2-tva-IRES-Cre* mice (referred to as *Olig2-Cre*) were crossed with *Rosa26-loxP-STOP-loxP-EYFP* (referred to as *Rosa26-EYFP*) reporter mice and the resulting double transgenic mice were analyzed at P17. **B**, low magnification confocal images of IHC for EYFP and Sox10 in the forebrain. Ctx, cerebral cortex, CC, corpus callosum, Hip, hippocampus. Boxed area was shown at higher magnification in **C**. Arrowheads, arrows, and wavy arrows point to EYFP^+^Sox10^+^ recombined OLCs, EYFP^-^Sox10^+^ OLCs escaping Cre-mediated recombination, and EYFP^+^Sox10^-^ cells, respectively. Scale bar = 20μm. **D**, strategy for disrupting PARP1 selectively in Olig2-expressing cells and their progeny cells. *Olig2-Cre* mice were crossed with *Parp1-exon4 floxed* mice (*Parp1^fl/fl^*) mice. The resulting mice of *Parp1*^fl/fl^, *Parp1*^fl/+^, or *Olig2-Cre* (collectively termed as *Parp1* Ctrl) and *Olig2-Cre:Parp1*^fl/fl^ (*Parp1* cKO) were analyzed at P14 (**E-H**) and adult (**I-M**) ages. **E-F**, representative low-magnification confocal images (**E**) and relative intensity (**F**) for myelin MBP and axonal SMI312 in the corpus callosum and overlying cingulate cortex of P14 mice. Scale bar = 200μm N = 3 Ctrl, 3 cKO, *t*_(4)_ = 2.839, *P* = 0.0469. **G-H**, representative confocal images of Sox10, PDGFRa, and CC1 (**G**) and quantification of Sox10^+^ OLCs, Sox10^+^PDGFRa^+^ OPCs, and Sox10^+^CC1^+^ OLs (**H**) in the corpus callosum of P14 mice. Scale bar = 10μm. N = 3 Ctrl, 3 cKO. OLCs *t*_(4)_ = 2.987, *P* = 0.0405; OPCs *t*_(4)_ = 2.645, *P* = 0.0573; OLs *t*_(4)_ = 3.027, *P* = 0.0389. **I-J**, representative images of Black Gold II myelin staining (**I**) and relative myelin intensity (**J**) in the corpus callosum and overlying cingulate cortex of P70 mice. Scale bar = 200μm. N = 5 Ctrl, 5 cKO, *t*_(8)_ = 0.3569, *P* = 0.7304. **K**, densities of Sox10^+^ OLCs, Sox10^+^PDGFRa^+^ OPCs, and Sox10^+^CC1^+^ OLs in the corpus callosum of P70 mice. N = 5 Ctrl, 5 cKO. OLCs *t*_(8)_ = 0.9632, *P* = 0.3637; OPCs *t*_(8)_ = 0.7271, *P* = 0.0.4878; OLs *t*_(8)_ = 1.225, *P* = 0.2554. **L**, accelerating Rotarod testing. Adult (P60) *Parp1* Ctrl and cKO mice were trained for 5 consecutive days (3 trials per day) followed by testing on day 6. Two-way ANOVA followed by Sidak multiple comparison test, genotype *F*_(1, 114)_ = 62.09 *P* < 0.0001; time-course *F*_(5, 114)_ = 22.74 *P* < 0.0001. M, *t*_(19)_ = 4.065, *P* = 0.0007. **M**, motor performance of *Parp1* Ctrl and cKO mice at Day 6, indicated by the latency time to fall. N=11 Ctrl, 10 cKO. *t*_(19)_ = 4.065, *P* = 0.0007 Statistic methods: unpaired two-tailed Student’s *t* test (F, H, J, K, M)) and two-way ANOVA followed by Sidak multiple comparison test (L). \**P* < 0.05; \*\**P* < 0.01; \*\*\**P* < 0.001, ns, not significant.

**fig. S4.**
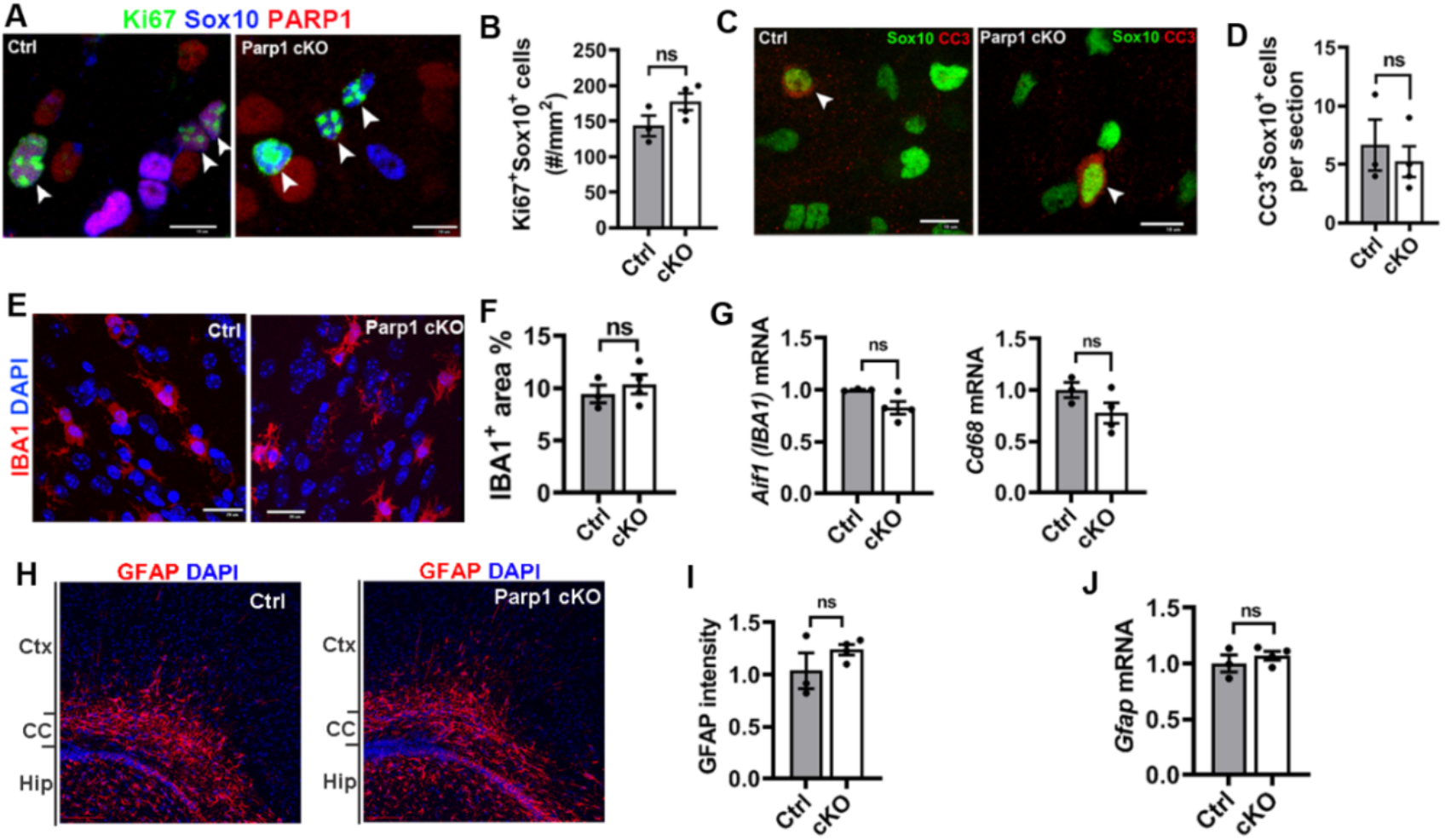
OPC-specific *Parp1* conditional knockout (cKO) does not perturb oligodendroglial proliferation, oligodendroglial cell death, and glial activation. *Parp* cKO (*Pdgfrα-CreER^T2^:Parp1^fl/fl^*) and Ctrl mice were i.p. injected with tamoxifen at P1, P2, and P3, and analyzed at P9. **A-B,** representative images (**A**) and density (**B**) of proliferating OPCs (Ki67^+^/Sox10^+^, arrowheads). N = 3 Ctrl and 4 cKO, *t*_(5)_ = 1.815, *P* = 0.1293. **C-D**, representative images (**C**) and quantification (**D**) of cleaved caspase 3 (CC3)^+^ apoptotic oligodendroglia. Arrowheads point to CC3^+^/Sox10^+^ dying cells. N = 3 Ctrl and 4 cKO. *t*_(5)_ = 0.5900, *P* = 0.5808. **E-F**, representative images of microglial marker IBA1 (**E**) and the percentage of IBA1-occupying area among total assessed area (**F**). Note that IBA1^+^ microglia exhibited similar morphology in *Parp1* Ctrl and cKO mice, both displaying “amoeboid-like” morphology as reported during CNS development. N = 3 Ctrl and 4 cKO. *t*_(5)_ = 0.7068, *P* = 0.5113. **G**, qRT-qPCR assay for *Iba1* and activated microglial marker *Cd68* in the brain. N = 3 Ctrl and 4 cKO. *Iba1*, Welch-corrected *t*_(2.792)_ = 3.507, *P* = 0.0668; *Cd68*, *t*_(5)_ = 1.686, *P* = 0.1527. **H-I**, representative confocal images (**H**) and quantification of astrocytic marker GFAP in the brain. N = 3 Ctrl and 4 cKO. *t*_(5)_ = 1.292, *P* = 0.2529. **J**, qRT-PCR assay for *Gfap* mRNA expression in the brain. N = 3 Ctrl and 4 cKO. *t*_(5)_ = 0.8999, *P* = 0.4094. Statistic methods: unpaired two-tailed Student’s *t* test, \**P* < 0.05; **P < 0.01; ***P < 0.001, ns, not significant. Panels A-F, H-I were collected from the corpus callosum and panels G and J from the whole brain of *Parp1* Ctrl and cKO mice. Scale bars: A, C, E, 10μm; H, 200μm.

**fig. S5.**
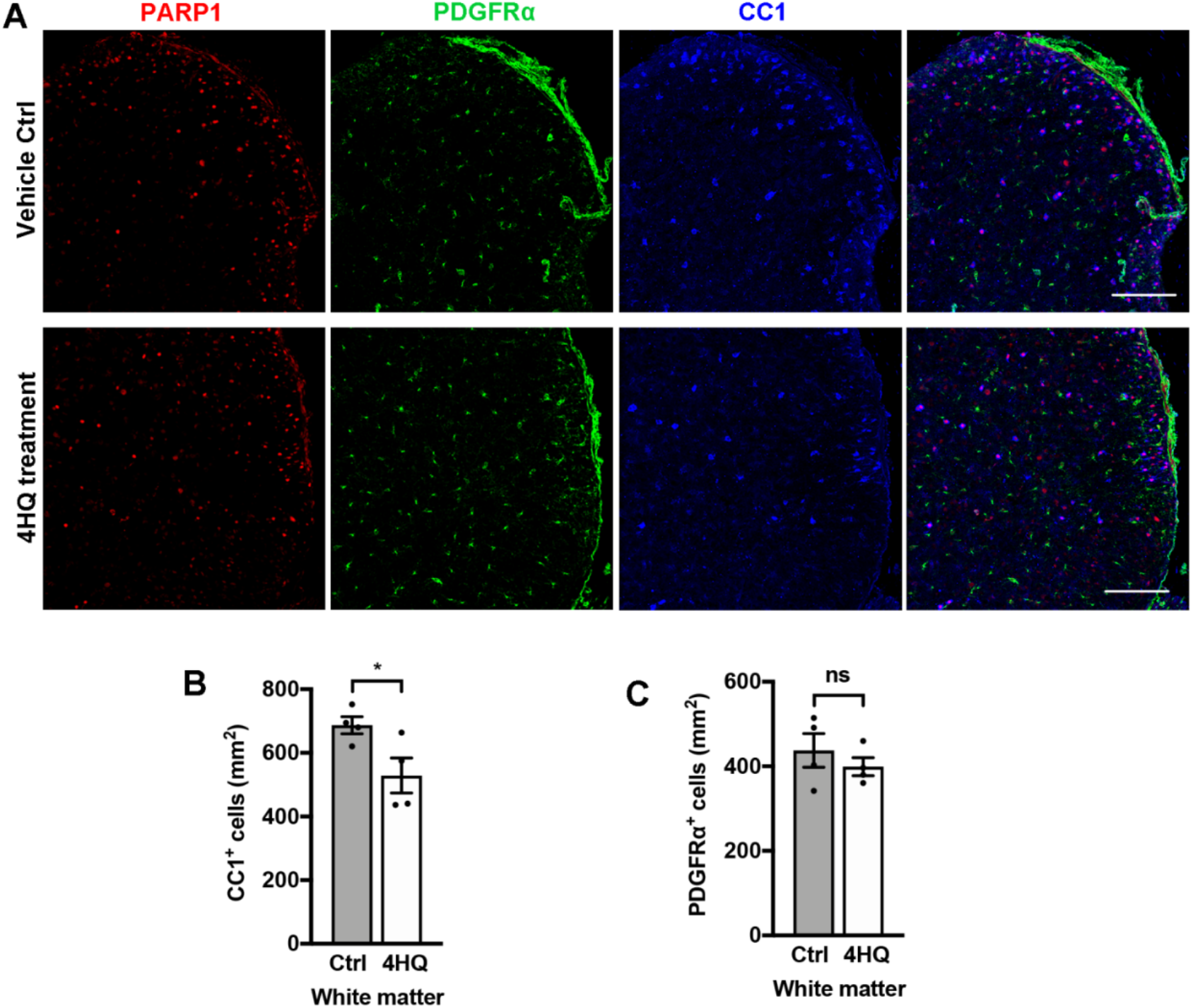
Inhibiting PARP1 activity by 4HQ perturbs OL formation in the spinal cord. Neonatal mice were injected with PARP1 inhibitor 4HQ daily at P2-P10 and analyzed 2 hr after the last injection at P10. **A**, representative confocal images of PARP1, PDGFRα, and CC1 in the spinal cord of 4HQ treated and vehicle treated control mice. Note that PARP1 expression is similar between Ctrl and 4HQ-treated spinal cord. Scale bar = 100μm. **B-C,** quantification of CC1^+^ OLs (**B**) and PDGFRα^+^ OPCs (**C**) in the white matter of spinal cord. N = 4 control, N = 4 4*HQ*. Two-tailed Student’s *t* test, N = 4 Ctrl and 4 4HQ. \**p* < 0.05; not significant *p* > 0.05. CC1^+^ OLs: *t*_(6)_ = 2.570, *P* = 0.0423; PDGFRα^+^ OPCs: *t*_(6)_ = 0.8507, *P* = 0.4276;

**fig. S6.**
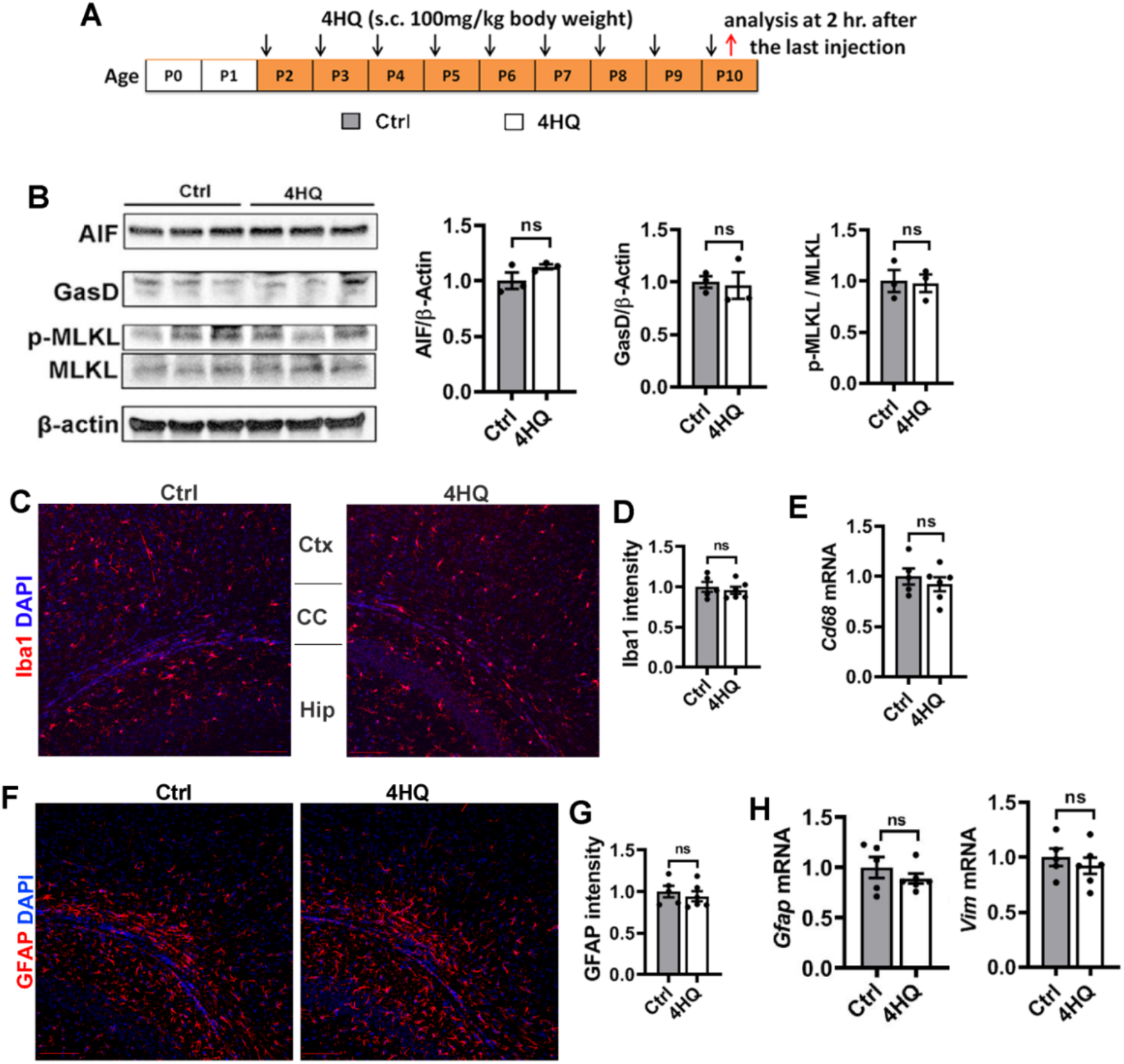
PARP1 inhibition by 4HQ does not affect cell death and glial activation. **A**, experimental design for panel **B-H**. **B**, Western blotting assay and quantification of forebrain protein for apoptosis-inducible factor (AIF, a mediator of PARP1-induced cell death under cytotoxic conditions), Gasdermin D (GasD, a mediator of pyroptotic cell death), and phosphorylated mixed lineage kinase domain-like protein (MLKL, Ser345), a trigger for necrotic cell death. N = 3 DMSO Ctrl, 3 4HQ. AIF *t*_(4)_ = 1.638, *P* = 0.1768; GasD *t*_(4)_ = 0.2424, *P* = 0.8204; p-MLKL *t*_(4)_ = 0.1618, *P* = 0.8793. **C-D**, representative confocal images (**C**) and relative intensity of Iba1 (**D**) in the forebrain. Ctx, cortex, CC, corpus callosum, Hip, hippocampus. N = 5 DMSO Ctrl, 6 4HQ, *t*_(9)_ = 0.5628, *P* = 0.5873. **E**, qRT-qPCR assay for activated microglial marker *Cd68* in the brain. N = 5 DMSO Ctrl, 6 4HQ, *t*_(9)_ = 0.7362, *P* = 0.4804.**F-G**, representative confocal images (**F**) and relative intensity of GFAP (**G**) in the forebrain. N = 5 DMSO Ctrl, 6 4HQ, *t*_(9)_ = 0.6155, *P* = 0.5535. **H**, qRT-qPCR assay for the mRNA levels of *Gfap* and activated astroglial marker Vimentin (*Vim*) in the brain. N = 5 DMSO Ctrl, 6 4HQ. *Gfap t*_(9)_ = 1.012, *P* = 0.3381; *Vim t*_(9)_ = 0.7043, *P* = 0.3381. Unpaired two-tailed Student’s *t* test, \**P* < 0.05; \*\**P* < 0.01; \*\*\**P* < 0.001; ns, not significant.

**fig. S7.**
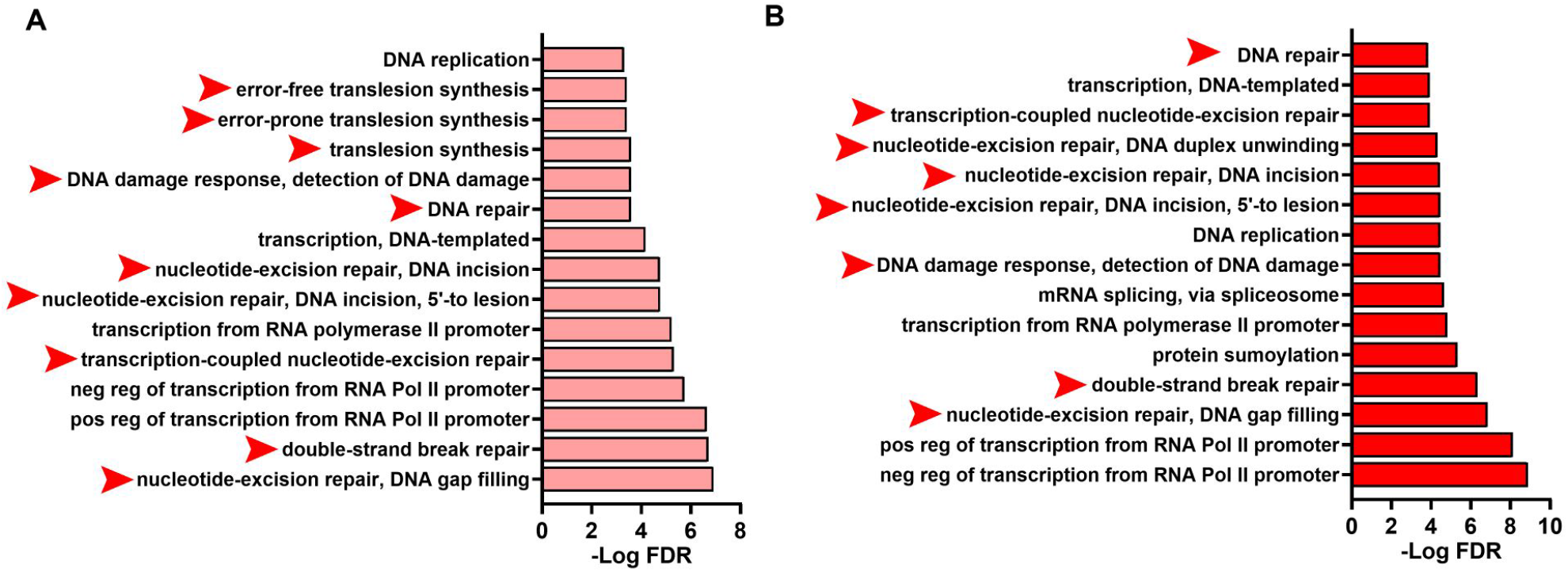
Top 15 GO terms of PARylated proteins identified in HCT116 cells injured by different concentration of DNA-damaging agent H_2_O_2_. (**A**, 0.2mM H_2_O_2_, **B**, 2mM H_2_O_2_). The identified PARylated proteins (129 proteins in **A** and 154 proteins in **B**) were retrieved from previous publication by Zhang et al., 2013. GO analysis were conducted using the Database for Annotation, Visualization and Integrated Discovery tool (DAVID, at https://david.ncifcrf.gov/tools.jsp). Note that biological processes related to DNA damage response and repair (red arrowheads) were significantly over-represented among the identified PARylated proteins at both H_2_O_2_ concentrations, which is consistent with the established role of PARP1 in DNA damage and repair.

## Notes

### Competing Interest Statement

The authors have declared no competing interest.

